# XACT-seq comprehensively defines the promoter-position and promoter-sequence determinants for initial-transcription pausing

**DOI:** 10.1101/2019.12.21.886044

**Authors:** Jared T. Winkelman, Chirangini Pukhrambam, Irina O. Vvedenskaya, Yuanchao Zhang, Deanne M. Taylor, Premal Shah, Richard H. Ebright, Bryce E. Nickels

**Author notes:** Equal contribution (Jared T. Winkelman, Chirangini Pukhrambam).

## Abstract

Pausing by RNA polymerase (RNAP) during transcription elongation, in which a translocating RNAP uses a “stepping” mechanism, has been studied extensively, but pausing by RNAP during initial transcription, in which a promoter-anchored RNAP uses a “scrunching” mechanism, has not. We report a method that directly defines RNAP-active-center position relative to DNA *in vivo* with single-nucleotide resolution (XACT-seq; crosslink-between-active-center-and-template sequencing). We apply this method to detect and quantify pausing in initial transcription at 4^11^ (∼4,000,000) promoter sequences *in vivo*, in *Escherichia coli*. The results show initial-transcription pausing can occur in each nucleotide addition during initial transcription, particularly the first 4-5 nucleotide additions. The results further show initial-transcription pausing occurs at sequences that resemble the consensus sequence element for transcription-elongation pausing. Our findings define the positional and sequence determinants for initial-transcription pausing and establish initial-transcription pausing is hard-coded by sequence elements similar to those for transcription-elongation pausing.

## Introduction

RNA polymerase (RNAP) initiates transcription by binding double-stranded promoter DNA, unwinding a turn of promoter DNA to yield an RNAP-promoter open complex (RPo) containing a ∼13 base pair single-stranded “transcription bubble” (Figure 1A) and selecting a transcription start site (TSS) (Ruff et al., 2015; Mazumder and Kapanidis, 2019; Winkelman et al., 2019). TSS selection entails placement of the start-site nucleotide (position +1) and the next nucleotide (position +2) of the template DNA strand into the RNAP-active-center product site (“P-site”) and addition site (“A-site”), respectively, and binding an initiating entity in the P-site and an extending NTP in the A-site.

**Figure 1.**
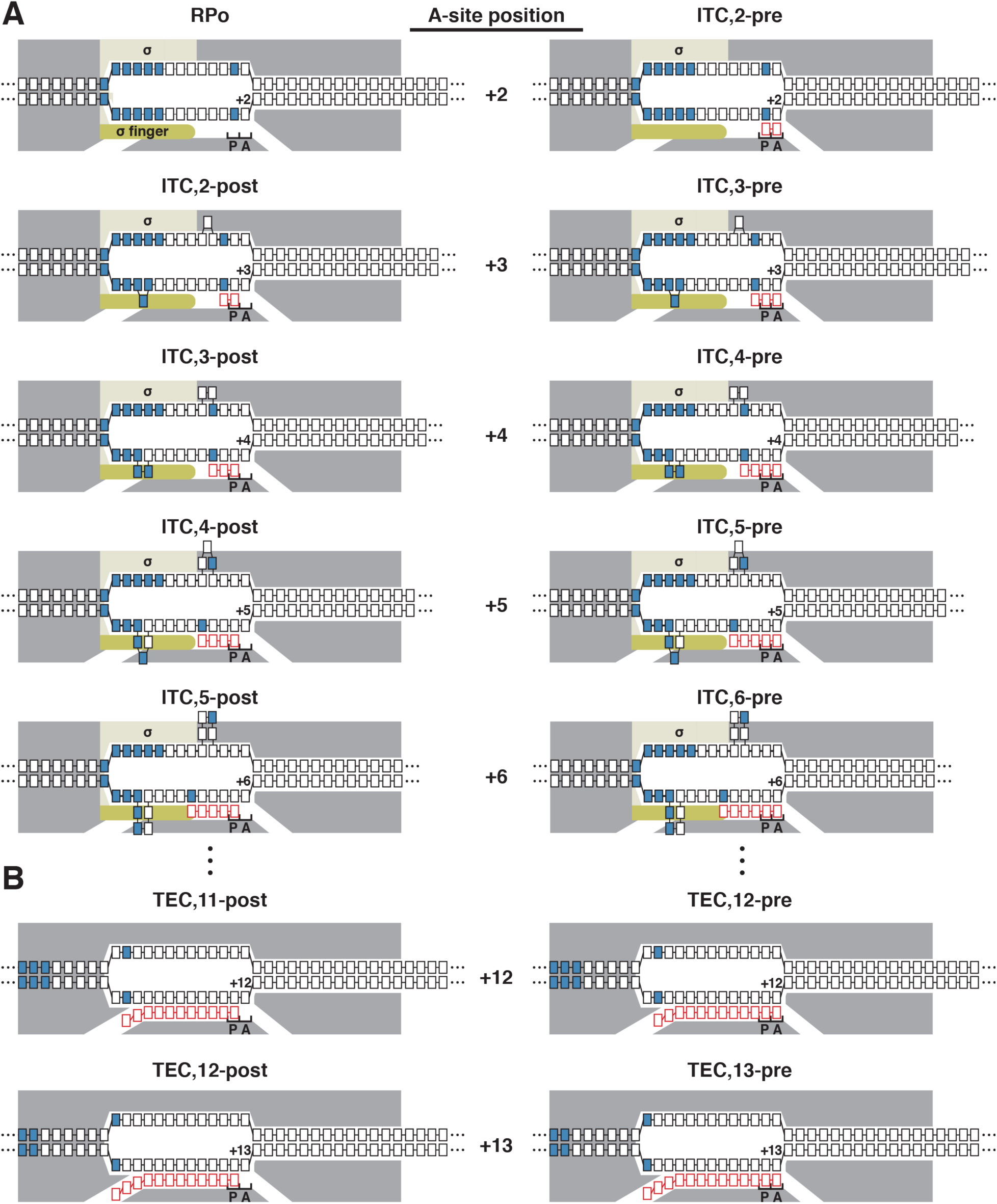
Mechanisms of initial transcription and transcription elongation. (A) Mechanism of initial transcription, with RNAP-active-center translocation through DNA scrunching mechanism and with RNAP in complex with transcription initiation factor σ. Panel shows first six nucleotide-addition steps of initial transcription, starting from the RNAP-promoter open complex (RPo), and yielding a series of successive RNAP-promoter initial transcribing complexes containing 2, 3, 4, 5, and 6 nt RNA products (ITC,2, ITC,3, ITC,4, ITC,5, and ITC,6). Following each nucleotide addition, the RNAP active center translocates forward through a DNA scrunching mechanism, from a pre-translocated state (pre; right column) to a post-translocated state (post; left column), and the RNAP-active-center A-site position advances by 1 bp (numbered positions between left and right columns). Grey, RNAP; yellow, σ; dark yellow, σ finger; blue, -10 element nucleotides and TSS nucleotides; P and A, RNAP-active-center P-site and A-site, respectively; black boxes, DNA nucleotides (nontemplate-strand nucleotides above template-strand nucleotides); red boxes, RNA nucleotides; promoter positions are numbered relative to the TSS position, +1. (B) Mechanism of transcription elongation, with RNAP-active center translocation through DNA stepping mechanism and with RNAP not containing σ. Panel shows two nucleotide-addition steps of transcription elongation, from a transcription elongation complex containing 11 nt of RNA (TEC,11) to a transcription elongation complex containing 12 nt of RNA (TEC,12). Following each nucleotide addition, RNAP, as a whole, translocates forward through a DNA stepping mechanism, between a pre-translocated state (pre; right column) and a post-translocated state (post; left column), and the RNAP-active-center A-site position advances by 1 bp (numbered positions between left and right columns). Symbols and colors as in panel A.

The first ∼10 nucleotides (nt) of an RNA product are synthesized as an RNAP-promoter initial transcribing complex (ITC) in which RNAP remains anchored on promoter DNA through sequence-specific protein-DNA interactions (Figure 1A; Mazumder and Kapanidis, 2019; Winkelman et al., 2019). Initial transcription starts with phosphodiester bond formation between the initiating entity and the extending NTP, to yield an initial RNA product (Figure 1A; ITC,2). Each nucleotide-addition cycle after formation of the initial RNA product requires translocation of the RNAP active center relative to DNA and RNA, starting from a “pre-translocated” state, and yielding a “post-translocated” state (Figure 1A,B; Erie et al., 1992; Zhang and Landick, 2009; Larson et al., 2011; Belogurov and Artsimovitch, 2019). Translocation of the RNAP active center repositions the RNA 3ʹ nucleotide from the RNAP-active-center “A-site” to the RNAP-active-center “P-site,” rendering the A-site available to bind the next extending NTP. Initial transcription proceeds until synthesis of an RNA product of a threshold length of ∼10 nt (Mazumder and Kapanidis, 2019; Winkelman et al., 2019). Upon synthesis of a threshold-length RNA product, RNAP breaks the sequence-specific protein-DNA interactions that anchor it on promoter DNA, escapes the promoter, and synthesizes the rest of the RNA product as a transcription elongation complex (TEC; Figure 1B).

There are two clear differences in the mechanism of initial transcription (performed by ITC) and transcription elongation (performed by TEC): (i) a different mechanism of RNAP active center translocation, and (ii) a different RNAP subunit composition (Figure 1).

The first difference in the mechanisms of initial transcription and transcription elongation is a consequence of the sequence-specific protein-DNA interactions that anchor RNAP on promoter DNA in initial transcription, but not in transcription elongation (Larson et al., 2011; Belogurov and Artsimovitch, 2019; Mazumder and Kapanidis, 2019; Winkelman et al., 2019). These sequence-specific protein-DNA interactions prevent RNAP from moving relative to DNA in initial transcription, but not in transcription elongation. Therefore, RNAP uses a different mechanism of RNAP active center translocation in initial transcription vs transcription elongation (Larson et al., 2011; Belogurov and Artsimovitch, 2019; Mazumder and Kapanidis, 2019; Winkelman et al., 2019). In initial transcription, RNAP uses a “scrunching” mechanism, in which, in each nucleotide-addition cycle, RNAP remains anchored to promoter DNA, unwinds one base pair of DNA downstream of the RNAP active center, pulls the unwound single-stranded DNA (ssDNA) into and past the RNAP active center, and accommodates the additional unwound ssDNA as bulges in the transcription bubble (Figure 1A; Kapanidis et al., 2006; Margeat et al., 2006; Revyakin et al., 2006). In contrast, in transcription elongation, RNAP uses a “stepping” mechanism, in which, in each nucleotide-addition cycle, RNAP steps forward by one base pair relative to the DNA (Figure 1B; Abbondanzieri et al., 2005).

The second difference in the mechanisms of initial transcription and transcription elongation is a consequence of the fact that initial transcription is carried out by a macromolecular assembly containing the initiation factor σ (Figure 1A), whereas transcription elongation is typically carried out by a macromolecular assembly lacking σ (Figure 1B; Mooney et al., 2005; Larson et al., 2011; Belogurov and Artsimovitch, 2019; Mazumder and Kapanidis, 2019; Winkelman et al., 2019). In the ITC, a module of σ referred to as the “σ finger,” reaches into the RNAP-active-center cleft and interacts with template-strand ssDNA in the transcription bubble close to the RNAP active center (Figure 1A; Severinov et al., 1994; Murakami et al., 2002; Kulbachinskiy and Mustaev, 2006; Zhang et al., 2012; Basu et al., 2014). These interactions pre-organize template-strand ssDNA to engage the RNAP active center, thereby facilitating the binding of initiating and extending NTPs (Murakami et al., 2002; Kulbachinskiy and Mustaev, 2006; Zhang et al., 2012; Pupov et al., 2014). Interactions of the σ finger with template-strand ssDNA places the σ finger in the path that will be occupied by nascent RNA upon RNA extension (Murakami et al., 2002; Kulbachinskiy and Mustaev, 2006; Zhang et al., 2012; Basu et al., 2014; Pupov et al., 2014; Zuo and Steitz, 2015). Thus, interactions of the σ finger with template-strand ssDNA presumably create an obstacle that must be overcome during initial transcription. Specifically, when the RNA reaches a length of ∼4 to ∼5 nt (after ∼2 to ∼3 RNAP-active-center translocation steps), the RNA 5ʹ end appears to contact, and make favorable interactions with, the σ finger, but, when the RNA reaches a length of ∼5 to ∼7 nt (after ∼3 to ∼5 RNAP-active-center translocation steps), the RNA 5ʹ end is expected to collide with, and clash with, the σ finger (Figure 1A; Murakami et al., 2002; Kulbachinskiy and Mustaev, 2006; Zhang et al., 2012; Basu et al., 2014; Pupov et al., 2014; Zuo and Steitz, 2015). The occurrence at one point of a favorable interaction with the σ finger (after ∼2 to ∼3 RNAP-active-center translocation steps), and subsequently a clash with the σ finger (after ∼3 to ∼5 RNAP-active-center translocation steps), is expected to have position-specific effects on RNAP active center translocation during initial transcription. In contrast, in transcription elongation, where σ typically is absent (Mooney et al., 2005; Belogurov and Artsimovitch, 2019), no such position-specific effects on RNAP-active-center translocation are expected to occur.

In both initial transcription and transcription elongation, in the presence of saturating NTPs, each nucleotide-addition cycle takes, on average, ∼20 ms (Belogurov and Artsimovitch, 2019). In both initial transcription and transcription elongation, pauses--nucleotide-addition cycles that occur on the second or longer timescales--are off-pathway states that potentially modulate gene-expression levels (Larson et al., 2011; Belogurov and Artsimovitch, 2019; Kang et al., 2019; Mazumder and Kapanidis, 2019). Transcription-elongation pausing has been the subject of extensive analysis, both *in vitro* and *in vivo* (Larson et al., 2011; Belogurov and Artsimovitch, 2019; Kang et al., 2019). The results establish that transcription-elongation pausing is determined by the sequence of the DNA template, rather than TEC position relative to the TSS or the length of RNA in the TEC. For *Escherichia coli* RNAP, both *in vitro* and *in vivo*, transcription-elongation pausing occurs at DNA sequence elements that have the consensus sequence G_-10_N_-9_N_-8_N_-7_N_-6_N_-5_N_-4_N_-3_N_-2_Y_-1_G_+1_, where Y is a pyrimidine and Y_-1_ corresponds to the position of the RNA 3’ end (Herbert et al., 2006; Larson et al., 2014; Vvedenskaya et al., 2014; Imashimizu et al., 2015; see also Kang et al., 2019; Saba et al., 2019). In contrast to transcription-elongation pausing, initial-transcription pausing has been subject to only limited studies of two promoters *in vitro* (Duchi et al., 2016; Lerner et al., 2016; Dulin et al., 2018) and no studies *in vivo*. As a result, it is not known whether initial-transcription pausing occurs *in vivo,* it is not known what fraction of promoter sequences undergo initial-transcription pausing, the promoter-position dependence of initial-transcription pausing has not been defined, and the promoter-sequence determinants for initial-transcription pausing have not been defined.

## RESULTS

### Rationale

During RNA synthesis, the dwell time of the RNAP active center at each transcribed-region position, “RNAP occupancy,” is correlated with the tendency of RNAP to pause at that position. Accordingly, pause sites can be identified using methods that provide a measure of RNAP occupancy (Figure 2). Previously reported sequencing-based methods to monitor RNAP occupancies, such as native elongating transcript sequencing (NET-seq), rely on identifying and quantifying RNA 3’ ends (Figure 2A; Churchman and Weissman, 2011; Churchman and Weissman, 2012; Larson et al., 2014; Vvedenskaya et al., 2014; Imashimizu et al., 2015). The identities of RNA 3’ ends allow estimation of RNAP-active-center positions--subject to uncertainties, due to ability of RNAP to sample pre-translocated, post-translocated, reverse-translocated, and hyper-translocated states (Figure 2A,B; Larson et al., 2011; Belogurov and Artsimovitch, 2019)--and the quantities of RNA 3’ ends allow estimation of RNAP-active-center dwell times. Unfortunately, however, these methods provide only an indirect measure of RNAP-active-center positions relative to DNA (Figure 2A), and because these methods cannot be applied to RNAs less than ∼15-nt in length (Figure S1), these methods are not suitable for analysis of initial transcription. Methods combining chromatin immunoprecipitation (ChIP) with sequencing, such as ChIP-seq and ChIP-exo, are alternative sequencing-based methods of mapping RNAP relative to DNA (Barski et al., 2007; Rhee and Pugh, 2012; Srivastava et al., 2013; Latif et al., 2018). However, these methods provide insufficient resolution for most purposes and define overall RNAP boundaries relative to DNA rather than RNAP-active-center A-site positions.

**Figure 2.**
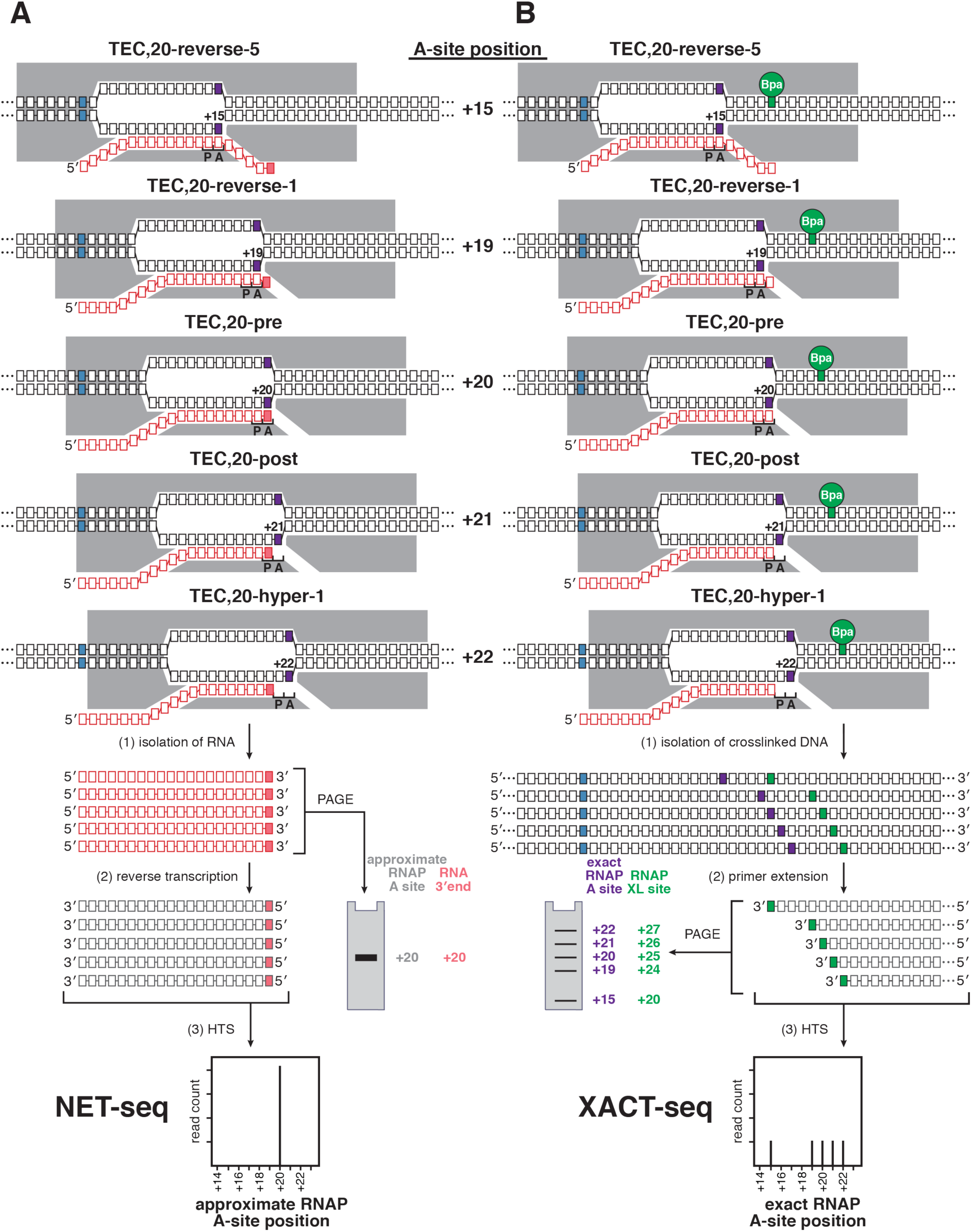
Approaches to map RNAP-active-center A-site position: NET-seq and XACT-seq. (A) Approach to map RNAP-active-center A-site position through analysis of RNA 3’ ends. The procedure entails isolating nascent RNA from transcription complexes and identifying and quantifying RNA 3’ ends by polyacrylamide gel electrophoresis (PAGE) or by analysis of primer extension products by high-throughput sequencing (HTS; NET-seq). The RNAP-active-center A-site position (purple boxes on nontemplate- and template-strand nucleotides; numbered relative to the TSS position, +1) of the promoter position in the RNAP-active-center A-site is approximated based on the identification of the RNA 3’ end (pink boxes); other symbols and colors as in Figure 1. The multiple different translocational states adopted by a transcription elongation complex (TEC) containing a single defined RNA product--e.g. reverse-translocated by 5 bp (TEC,20-reverse-5), reverse-translocated by 1 bp (TEC,20-reverse-1), pre-translocated (TEC, 20-pre), post-translocated (TEC,20-post), and hyper-translocated by 1 bp (TEC,20-hyper-1)--all yield the same RNA 3’ end and therefore cannot be distinguished. (B) Approach to map the RNAP-active-center A-site position by site-specific protein-DNA photo-crosslinking with an RNAP derivative that contains a photo-activatable agent (green circle labeled Bpa) that crosslinks to DNA a defined distance from the RNAP active center. The procedure entails UV irradiating transcription complexes, isolating RNAP-crosslinked DNA, and identifying and quantifying crosslinking sites by analysis of primer extension products by PAGE or HTS (XACT-seq). The RNAP-active-center A-site position (purple boxes on template and nontemplate strands; numbered relative to the TSS position, +1) is defined based on the identity of the crosslinking site (green boxes; XL site); other symbols as in panel A. The multiple different translocational states adopted by a transcription elongation complex (TEC) containing a single defined RNA product--e.g. reverse-translocated by 5 bp (TEC,20-reverse-5), reverse translocated by 1 bp (TEC,20-reverse-1), pre-translocated (TEC, 20-pre), post-translocated (TEC,20-post), and a hyper-translocated by 1 bp state (TEC,20-hyper-1)--all yield different crosslinking sites and therefore can be distinguished.

Here we describe a sequencing-based method to monitor RNAP occupancy that overcomes these limitations, providing a direct, single-nucleotide-resolution readout of RNAP-active-center position relative to DNA (Figure 2B), and that is suitable for analysis of initial transcription as well as transcription elongation. The method entails formation of transcription complexes *in vitro* or *in vivo* using an RNAP derivative that has a photo-activatable crosslinking agent incorporated at a single, defined site in RNAP that, upon photo-activation *in vitro* or *in vivo*, forms covalent crosslinks with DNA at a defined position relative to the RNAP-active-center A-site; photo-activation to initiate covalent crosslinking of RNAP to DNA *in vitro* or *in vivo*; and high-throughput sequencing of primer extension products to define crosslink positions and crosslink yields. We term this method: “<u>cross</u>link-between-<u>a</u>ctive-<u>c</u>enter-and-template sequencing” (XACT-seq; Figure 2B).

XACT-seq takes advantage of an RNAP derivative that has a photo-activatable crosslinking amino acid *p*-benzoyl-L-phenylalanine (Bpa) incorporated at RNAP-β’ subunit residue R1148 (RNAP-β′^R1148Bpa^), which, upon photo-activation *in vitro* or *in vivo*, forms covalent crosslinks with DNA at a position exactly 5-nt downstream of the RNAP-active-center A-site (Figure 2B; Yu et al., 2017). In previous work, this reagent was used for structural analysis of static, trapped transcription complexes (Winkelman et al., 2015; Winkelman et al., 2016a; Yu et al., 2017). In this work, we apply the reagent for analysis of actively-transcribing complexes.

### RNAP-active-center A-site positions in initial-transcription pausing at the *lac*CONS promoter *in vitro* and *in vivo*

To demonstrate that the RNAP derivative that underpins XACT-seq, RNAP-β′^R1148Bpa^, enables detection of RNAP-active-center A-site positions in actively-transcribing complexes, and enables detection of initial-transcription pausing, we used RNAP-β′^R1148Bpa^ to analyze actively-transcribing complexes engaged in initial-transcription pausing (Figure 3). We analyzed initial-transcription pausing at the *lac*CONS promoter (p*lac*CONS; Figure 3A, top), the best-characterized example of initial-transcription pausing (Duchi et al., 2016; Lerner et al., 2016; Mazumder and Kapanidis, 2019). Lerner *et al*. and Duchi *et al*. showed *in vitro* that initial-transcription pausing occurs at p*lac*CONS after ∼4 to ∼6 RNAP-active-center translocation steps (Duchi et al., 2016; Lerner et al., 2016). Duchi *et al*. and Dulin *et al*. further showed *in vitro* that this pause is reduced upon deletion of the σ finger (Duchi et al., 2016) and is increased upon substitution of the RNAP β-subunit residue 446 (β^D446A^) (Dulin et al., 2018), a substitution that alters the sequence dependence of RNAP-active-center translocation behavior (Zhang et al., 2012; Vvedenskaya et al., 2014).

**Figure 3.**
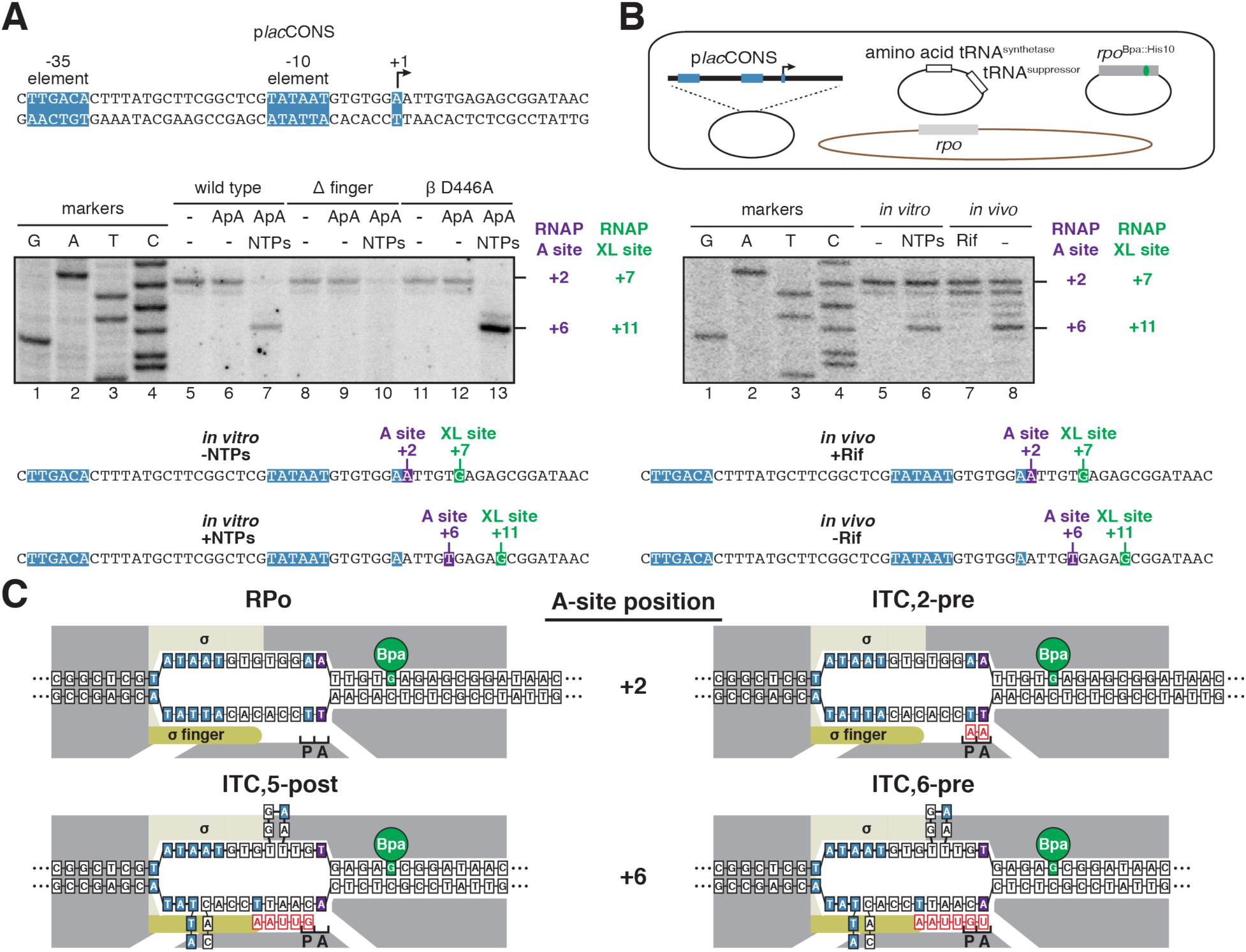
RNAP-active-center A-site positions in initial-transcription pausing at the *lac*CONS promoter *in vitro* and *in vivo*. (A) Analysis of initial-transcription pausing at p*lac*CONS by RNAP-β′^R1148Bpa^ *in vitro*. Top, *lac*CONS promoter (p*lac*CONS). Middle, results. Bottom, position of RNAP-active-center A-site (purple) and nucleotide crosslinked to Bpa (green) defined relative to the TSS position. Markers, DNA sequence ladder generated using p*lac*CONS template. (B) Analysis of initial-transcription pausing at p*lac*CONS by RNAP-β′^R1148Bpa^ *in vivo*. Top, three-plasmid merodiploid system for co-production, in *E. coli* cells, of decahistidine-tagged, RNAP-β′^R1148Bpa^, in the presence of untagged wild-type RNAP. The first plasmid carries gene for RNAP subunit with a nonsense (TAG) codon at residue βʹ R1148 (grey rectangle; green indicates nonsense codon); The second plasmid carries genes for engineered Bpa-specific nonsense-supressor tRNA and Bpa-specific amino-acyl-tRNA synthetase (white rectangles); the third plasmid carries p*lac*CONS; and the chromosome (brown oval) carries wild-type RNAP subunit genes (light grey rectangle). Middle, results. Bottom, position of RNAP-active-center A-site (purple) and nucleotide crosslinked to Bpa (green) defined relative to the TSS position. Rif, Rifampin; Markers, DNA sequence ladder generated using p*lac*CONS template. (C) Interpretation of results in (A) and (B). Symbols and colors as in Figures 1,2.

We first used RNAP-β′^R1148Bpa^ to detect initial-transcription pausing at p*lac*CONS *in vitro* (Figure 3A). The results in Figure 3A, lanes 5, 8, and 11, confirm the previously reported ability of the RNAP-β′^R1148Bpa^ to detect the RNAP-active-center A-site position with single-nucleotide resolution in static initial-transcribing complexes in the absence of NTP substrates *in vitro* (Winkelman et al., 2015; Winkelman et al., 2016a; Winkelman et al., 2016b; Yu et al., 2017). The results in Figure 3A, lanes 6, 9, and 12, demonstrate that the detected position of the RNAP-active-center A-site position does not change upon addition of a 2-nt RNA (ApA), indicating that the resulting ITC,2 is in a pre-translocated state (ITC,2-pre). The results in Figure 3A, lane 7, demonstrate the ability of RNAP-β′^R1148Bpa^ to detect RNAP-active-center A-site positions--and to detect initial-transcription pausing--in active initial-transcribing complexes in the presence of all NTP substrates *in vitro*. We observe pausing in the initial-transcribed region (positions +3 to +9; Figure 3) and in the downstream transcribed-region sequence (positions ≥ +10). For the purpose of this paper we focus on the initial-transcribed region. Results for promoter occupancies during transcription elongation will be reported elsewhere.

We observe strong pausing at a position corresponding to exactly 4 RNAP-active-center translocation steps relative to RPitc,2-pre (i.e., when the RNAP-active-center A-site is at promoter position +6; Figure 3A). We observe that RNAP occupancy at this position decreases upon deletion of the σ finger (Figure 3A; compare lanes 7 and 10) and increases upon substitution of RNAP β-subunit residue 446 (Figure 3A; compare lanes 7 and 13). This pause exhibits the previously reported hallmarks of initial-transcription pausing at p*lac*CONS: i.e., pausing after ∼4 to ∼6 active-center translocation steps (Duchi et al., 2016; Lerner et al., 2016), a decrease in pausing upon deletion of the σ finger (Duchi et al., 2016), and an increase in pausing upon substitution of RNAP β-subunit residue 446 (Dulin et al., 2018). We conclude that the pausing that occurs when the RNAP-active-center A-site is at promoter position +6 represents initial-transcription pausing.

We next used RNAP-β′^R1148Bpa^ to detect initial-transcription pausing at p*lac*CONS *in vivo* (Figure 3B). We produced, in *Escherichia coli*, a Bpa-labeled, decahistidine-tagged RNAP-β′^R1148Bpa^ in the presence of unlabeled, untagged, wild-type RNAP, using a three-plasmid system (Figure 3B, top) comprising (i) a plasmid carrying a gene for RNAP β’-subunit containing a UAG codon at position 1148 and a decahistidine coding sequence; (ii) a plasmid carrying genes for an engineered Bpa-specific UAG-suppressor tRNA and an engineered Bpa-specific aminoacyl-tRNA synthetase; and (iii) a plasmid containing p*lac*CONS. We then grew cells in medium containing Bpa, UV irradiated cells, lysed cells, purified crosslinked material using immobilized metal-ion-affinity chromatography (IMAC) targeting the decahistidine tag on RNAP-β′^R1148Bpa^, and mapped crosslinks using primer extension (Figure 3B, middle). In order to trap static initial-transcribing complexes containing 2- to 3-nt RNA products *in vivo*, despite the presence of all NTP substrates, we used a chemical biology approach, exploiting the RNAP inhibitor Rifampin (Rif), which blocks extension of RNA products beyond a length of 2 to 3 nt (McClure and Cech, 1978). The results showed exact correspondence of crosslinking patterns *in vitro* and *in vivo* for the initial-transcription pause at promoter position +6 (Figure 3B).

The results in Figure 3 establish that RNAP-β′^R1148Bpa^ enables detection of the RNAP-active-center A-site position in actively-transcribing complexes. The results further establish that initial-transcription pausing at p*lac*CONS occurs *in vivo* and exhibits the same promoter-position dependence *in vivo* as *in vitro*. The results reveal the exact RNAP-active-center position in initial-transcription pausing at p*lac*CONS. In particular, the results show that the RNAP-active-center A-site during initial-transcription pausing at p*lac*CONS is located at promoter position +6 and thus, neglecting possible fractionally-translocated states, show that pausing occurs in either RPitc,5-post or RPitc,6-pre (Figure 3C).

### Promoter-position dependence of initial-transcription pausing for a library of 4^11^ (∼4,000,000) promoters *in vivo*

Having validated RNAP-β′^R1148Bpa^ as a reagent for detection of the RNAP-active-center A-site position in actively-transcribing complexes *in vitro* and *in vivo*, we next combined RNAP-β′^R1148Bpa^ with a library of p*lac*CONS derivatives (p*lac*CONS-N11; Figure 4A) containing all possible sequences from promoter position +3 to promoter position +13 (4^11^; ∼4,000,000 sequences), in order to enable multiplexed analysis of initial-transcription pausing on all possible initial-transcribed region sequences *in vivo* (Figures 4, 5, S2, S3). We used a three-plasmid system analogous to that used in the previous section but having representatives of the p*lac*CONS-N11 library of plasmids instead of the plasmid carrying p*lac*CONS. We UV irradiated cells to initiate RNAP-DNA crosslinking, lysed cells, isolated crosslinked material, and mapped crosslinks using primer extension as in the previous section. We used the same chemical-biology approach, exploiting the fact that the RNAP inhibitor Rif prevents extension of RNA products beyond the length of 2 to 3 nt to trap static initial-transcribing complexes corresponding to RPitc,2 and RPitc,3 (Figure 4B).

**Figure 4.**
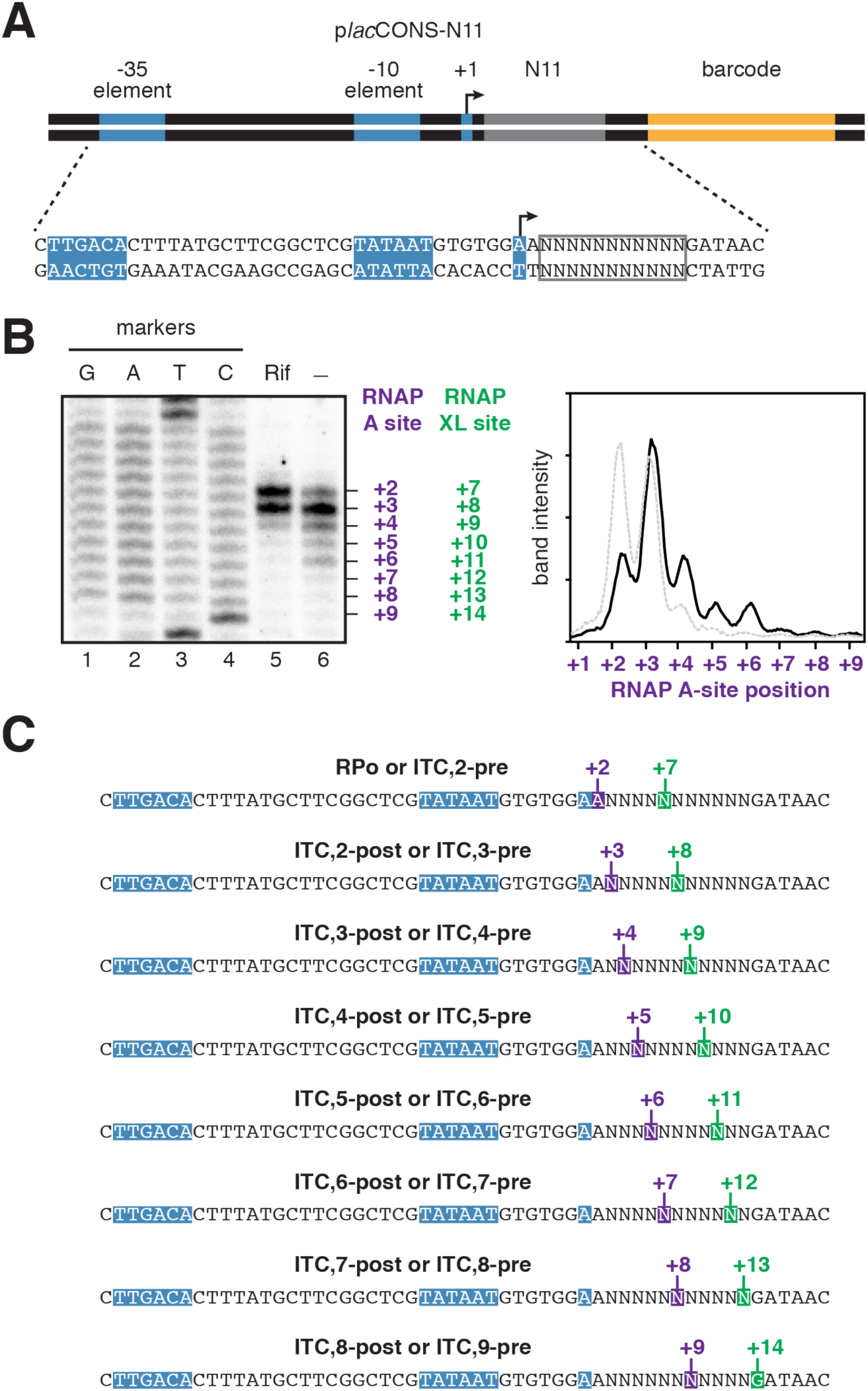
Promoter-position dependence of initial-transcription pausing for a library of 4^11^ (∼4,000,000) promoters *in vivo*. (A) p*lac*CONS template library containing all possible sequences from promoter position +3 to promoter position +13 (p*lac*CONS-N11). (B) Left, PAGE analysis of RNAP-active-center A-site positions *in vivo*. Right, histogram showing signals detected in the absence (black line) or presence (grey line) of Rif. Markers, DNA sequence ladder generated using p*lac*CONS-N11. (C) Position of RNAP-active-center A-site (purple) and nucleotide crosslinked to Bpa (green) defined relative to the TSS position, +1.

In order to read out position-specific RNAP occupancies across the entire sampled initial-transcribed-region sequence space, we performed urea-PAGE of radiolabeled primer extension products (Figure 4B). In the presence of Rif, we see two major bands and a minor band across the sampled initial-transcribed-region sequence space at positions corresponding to RPitc,2; RPitc,3; and RPitc,4--i.e. when the RNAP-active-center A-site is at promoter positions +2, +3, or +4 (see Figure 1A)--exactly as above for the p*lac*CONS initial-transcribed-region sequence (compare Figure 4B, lane 5 with Figure 3B, lane 7). In the absence of Rif, we observe a series of additional bands corresponding to positions of initial-transcription pausing (Figure 4B, lane 6). The prominence of the additional bands indicates that initial-transcription pausing is a prominent feature of transcription across the entire sampled initial-transcribed-region sequence space. As compared to the experiments in the presence of Rif, in the absence of Rif, we observe higher levels of RNAP occupancy at positions corresponding to exactly 1, 2, 3, 4, 5, 6, and 7 RNAP-active-center translocation steps relative to RPitc,2-pre (i.e., when the RNAP-active-center A-site is at promoter positions +3, +4, +5, +6, +7, +8, and +9; Figure 4B,C). RNAP occupancy levels are highest at the positions corresponding to 1 to 4 RNAP-active-center translocation steps (i.e., when the RNAP-active-center A-site is at promoter positions +3, +4, +5, and +6; Figure 4B,C) and are lower for the positions corresponding to 5, 6, or 7 RNAP-active-center translocation steps (i.e., when the RNAP-active-center A-site is at promoter positions +7, +8, and +9; Figure 4B,C). Similar results are observed in parallel experiments using a promoter library that randomizes 20 bp of the initial-transcribed region, from position +2 through +21 (Figure S2). We conclude that, across the sampled initial-transcribed-region sequence space, initial-transcription pausing occurs at a large fraction of promoters. We further conclude that, across the sampled initial-transcribed-region sequence space, initial-transcription pausing can occur at any promoter initial-transcribed-region position from position +3 through position +9, with highest levels for promoter positions +3 through +6.

### Promoter-sequence determinants for initial-transcription pausing in a library of 4^11^ (∼4,000,000) promoters *in vivo*

In order to quantify the promoter position dependence of initial-transcription pausing, and in order to define the promoter sequence determinants for initial-transcription pausing, for the library of 4^11^ (∼4,000,000) initial-transcribed-region sequences, we performed the full XACT-seq protocol (Figure S3). We used the same three-plasmid system and the same procedure of UV irradiation, cell lysis, purification of crosslinked material, and primer extension as in the previous sections. We then used high-throughput sequencing to identify and quantify primer extension products. We identified RNAP-active-center A-site positions based on the identities of 3’ ends of primer extension products, we quantified RNAP occupancies at these positions based on numbers of sequence reads, and we assigned sites of initial-transcription pausing as sites with above-threshold RNAP occupancies.

The results confirm and quantify the finding in the preceding section that, across the sampled initial-transcribed-region sequence space, a substantial fraction of promoter initial-transcribed region sequences--∼20% of promoter sequences--show initial-transcription pausing (Table S1). The results further confirm and quantify the finding in the preceding section that, across the sampled initial-transcribed-region sequence space, initial-transcription pausing can occur at each promoter initial-transcribed-region position, from position +3 through position +9, with highest levels occurring when the RNAP-active-center A-site is at positions +3 through +7, and lower levels when the RNAP-active-center A-site is at positions +8 and +9 (Table S1).

A global alignment of all initial-transcribed-region sequences showing initial-transcription pausing--aligning by the RNAP-active-center A-site position--yields a clear consensus sequence: T_P-1_N_P_<u>Y</u>_A_G_A+1_, where Y is pyrimidine and <u>Y</u>_A_ is the position of the RNAP-active-center A-site (Figure 5A,B). A plot of RNAP occupancy as a function of initial-transcribed-region tetranucleotride sequence shows graphically that RNAP occupancy is strongly correlated with match to the consensus sequence (Figure 5B). Separate alignments of initial-transcribed-region sequences showing pausing for each promoter position from +3 through +9, yield, in all cases, a consensus sequence resembling the global alignment (Figures 5C, S4). Plots of RNAP occupancy as a function of initial-transcribed-region sequence shows graphically that RNAP occupancy strongly correlates with match to the consensus sequence for each position (Figure S4). We note that p*lac*CONS, the promoter previously shown to exhibit initial-transcription pausing at a position in the range +5 to +7 *in vitro* (Duchi et al., 2016; Lerner et al., 2016; Dulin et al., 2018), and shown here to exhibit initial transcription pausing at position +6 *in vitro* and *in vivo* (Figure 3), contains a 3-of-3 match to the consensus sequence in the register that would yield initial transcription pausing at position +6 (compare Figure 3A and Figure 5A).

**Figure 5.**
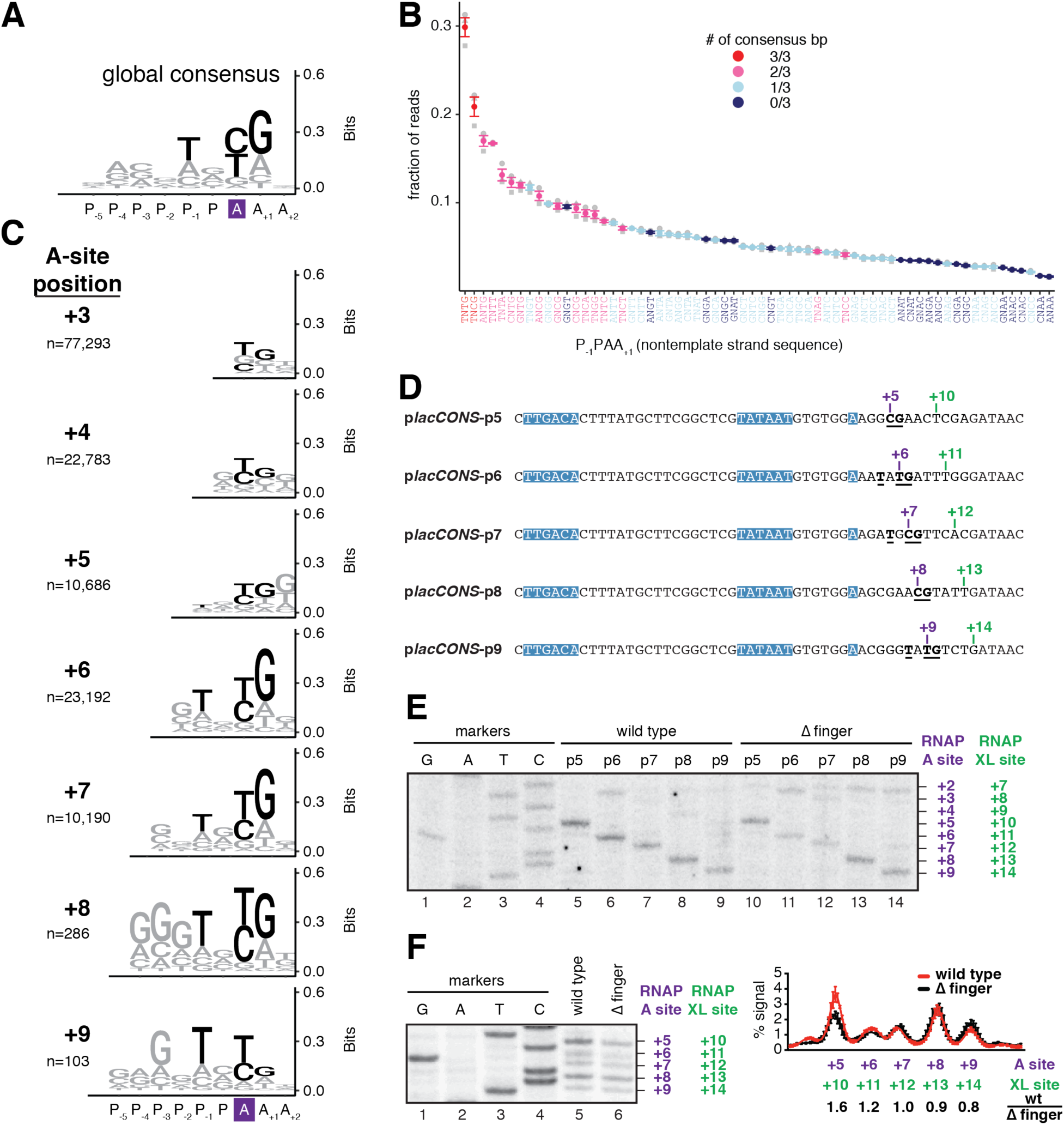
Promoter-sequence determinants for initial-transcription pausing in a library of 4^11^ (∼4,000,000) promoters *in vivo*. (A) Position-independent, global, consensus sequence for initial-transcription pausing (B) RNAP occupancy at each initial-transcribed region tetranucleotide sequence. Red, pink, cyan, and blue denote pause-site sequences with 3 of 3, 2 of 3, 1 of 3, and 0 of 3 matches to the global consensus sequence. Mean ± SEM (n = 3). (C) Position-specific consensus sequences for initial-transcription pausing for RNAP-active-center A-site positions +3, +4, +5, +6, +7, +8, +9. (D) Representative sequences yielding high RNAP occupancy at RNAP-active-center A-site positions +5, +6, +7, +8, +9, respectively. Colors as in Figure 4C. (E) RNAP-active-center A-site positions and crosslinking positions *in vitro* for the sequences of (C) with high RNAP occupancy at RNAP-active-center A-site positions +5, +6, +7, +8, +9. Lanes 1-4 present sequence markers, lanes 5-9 present data for wild-type RNAP, and lanes 10-14 present data for an RNAP derivative containing a deletion of the σ finger (Δ finger). (F) RNAP-active-center A-site positions and crosslinking positions *in vitro* in which sequences of (C) are pooled and analyzed as a pool. Left, gel; right subpanel presents quantitation of RNAP occupancies at +5, +6, +7, +8 and +9, with data for wild-type RNAP in red and data for Δ finger RNAP in black.

Representative individual initial-transcribed-region sequences showing initial-transcription pausing were analyzed for pausing individually, both *in vitro* and *in vivo*, and showed clear pausing at the expected positions *in vitro* and *in vivo* (Figures 5D-F, S5). The staircase pattern in Figure 5E (*in vitro*) and Figure S5B (*in vivo*) shows graphically that initial-transcription pausing can occur at each promoter position in the tested range.

We performed analogous experiments with individual initial-transcribed-region sequences using an RNAP holoenzyme derivative containing a deletion of the σ finger (Figure 5E,F). Qualitatively, the results for wild-type RNAP holoenzyme and the RNAP holoenzyme derivative containing the σ finger deletion were identical for individual initial-transcribed-region sequences (Figure 5E,F), indicating that sequence, rather than position-dependent collision with the σ finger, is the crucial determinant for initial-transcription pausing. Quantitatively, however, we observe reductions in RNAP occupancies for the RNAP holoenzyme containing the σ finger, relative to wild-type RNAP holoenzyme, for promoter positions +5 and +6, but not for promoter positions +7, +8 and +9 (Figure 5E,F). We infer that the σ finger contributes quantitatively to pausing at promoter positions +5 and +6, which are positions where collision of the RNA 5ʹ end and the σ finger potentially may occur (Murakami et al., 2002; Kulbachinskiy and Mustaev, 2006; Zhang et al., 2012; Basu et al., 2014; Pupov et al., 2014; Zuo and Steitz, 2015).

Considering the results in Figure 5, as a whole, we conclude that sequence is the crucial determinant of initial-transcription pausing. We further conclude that sequence is the crucial determinant of initial-transcription pausing at each promoter position from position +3 to position +9, and that the consensus sequences for initial-transcription pausing is similar or identical at each position.

### Relationship between sequence determinants for initial-transcription pausing and sequence determinants for transcription-elongation pausing

Our results define the consensus sequence for initial-transcription pausing as: T_P-1_N_P_<u>Y</u>_A_G_A+1_ (Figure 6A, top). This is both the global consensus sequence for initial-transcription pausing irrespective of position and, in most, and possibly all, cases the position-specific sequence for initial-transcription pausing at each initial-transcribed region position (Figure 5A-C).

**Figure 6.**
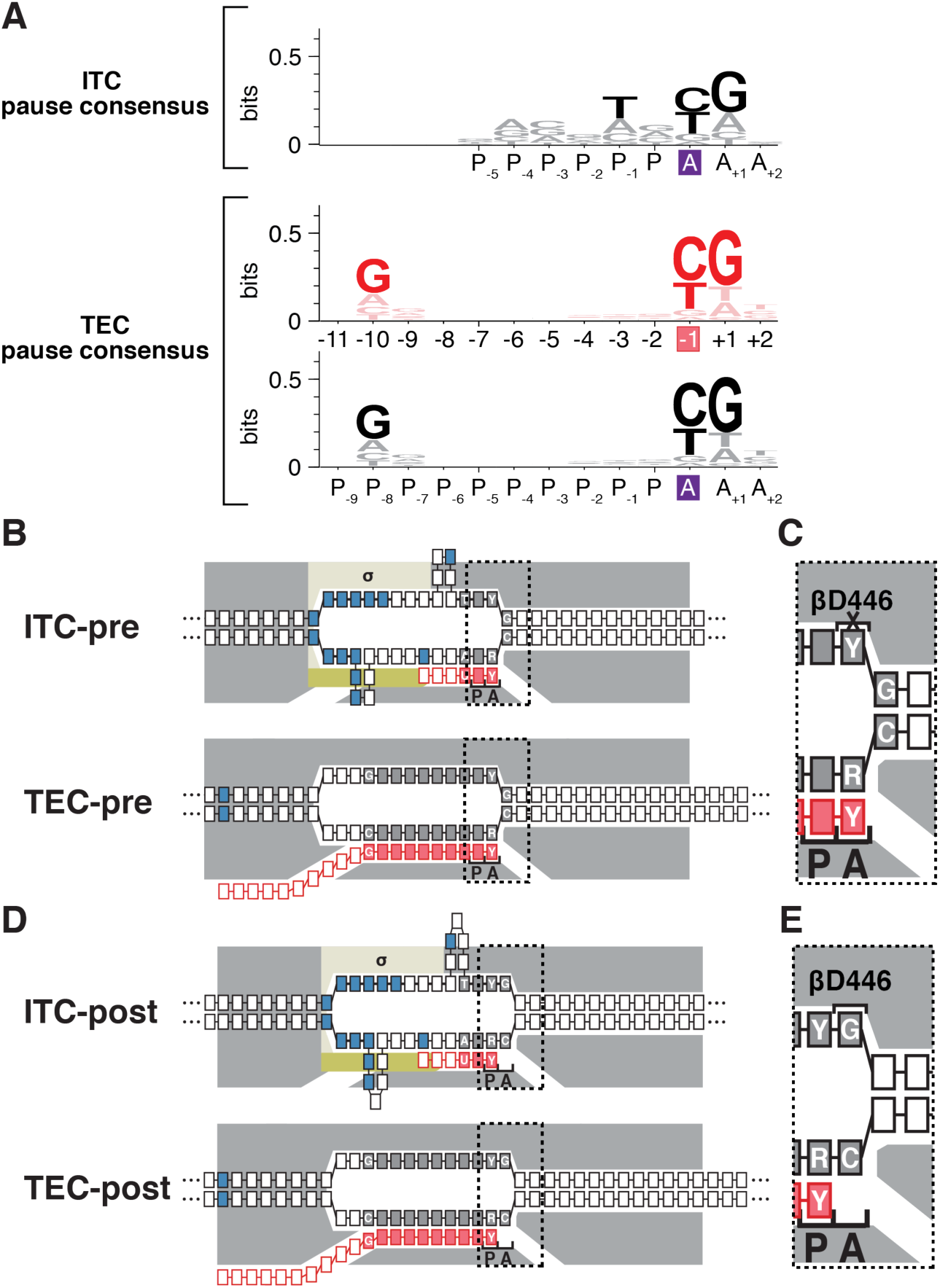
Relationship between sequence determinants for initial-transcription pausing and sequence determinants for transcription-elongation pausing. (A) Consensus sequence for initial-transcription pausing. Positions are labeled relative to RNAP-active-center A-site (purple box) and P-site. (B) Consensus sequence for transcription-elongation pausing. Positions are numbered relative to the RNA 3’ end (pink box) in upper axis label and numbered relative to RNAP-active-center A-site (purple box) and P-site in lower axis label. (C) Positions of consensus-sequence nucleotides in pre-translocated-state complexes in initial-transcription pausing (ITC-pre) and transcription elongation pausing (TEC-pre). Grey boxes with white lettering, high-information content DNA nucleotides of consensus sequence; grey boxes without white lettering; low-information content nucleotides of consensus-sequence; Pink boxes with white lettering, high-information content RNA nucleotides of consensus sequence. Dotted rectangle, region enlarged at right. Other symbols and colors as in Figure 1. (D) Enlarged view of RNAP-active-center region of pre-translocated state complexes, showing unfavorable interaction of RNAP β residue D446 with nontemplate-strand pyrimidine (Y) at A-site (crossed out bracket labeled β-D446). (E) Positions of consensus-sequence nucleotides in post-translocated-state complexes in initial transcription pausing (ITC-post) and transcription elongation pausing (TEC-post). (F) Enlarged view of RNAP-active-center region of post-translocated state complexes, showing favorable interaction of RNAP β residue D446 with nontemplate-strand purine (R) at A-site (bracket labeled β-D446).

Three groups previously have used NET-seq to define the consensus sequence for transcription-elongation pausing: G_-10_N_-9_N_-8_N_-7_N_-6_N_-5_N_-4_N_-3_N_-2_Y_-1_G_+1_, where Y_-1_ is the position of the RNA 3ʹ end (Figure 6A, middle; Larson et al., 2014; Vvedenskaya et al., 2014; Imashimizu et al., 2015). Mapping of TEC translocational state by analysis of pyrophosphorolysis kinetics (Vvedenskaya et al., 2014) and by cryo-EM structure determination (Guo et al., 2018; Kang et al., 2018) indicates that TECs paused at the consensus sequence element for transcription-elongation pausing assume a pre-translocated or fractionally-translocated state. Accordingly, the consensus sequence for transcription-elongation pausing can be expressed as: G_P-8_N_P-7_N_P-6_N_P-5_N_P-4_N_P-3_N_P-2_N_P-1_N_P_<u>Y</u>_A_G_A+1_, where <u>Y</u>_A_ is the position of the RNAP-active-center A-site (Figure 6A, bottom). The consensus sequence for transcription elongation pausing, G_P-8_N_P-7_N_P-6_N_P-5_N_P-4_N_P-3_N_P-2_N_P-1_N_P_<u>Y</u>_A_G_A+1_, comprises a highly-conserved downstream segment, <u>Y</u>_A_G_A+1_ (Larson et al., 2014; Vvedenskaya et al., 2014; Imashimizu et al., 2015). In the paused TEC, the G_A+1_:C_A+1_ nucleotide pair is located at the downstream end of the transcription bubble and must be broken for RNAP forward translocation, and the DNA-template-strand <u>R</u>_A_ interacts with the RNAP active center (Figure 6B,C; Vvedenskaya et al., 2014; see also Saba et al., 2019); the identities of nucleotides at these positions affects RNAP-translocation behavior through sequence-dependent effects on DNA duplex thermal stability and sequence-dependent effects on interactions of the DNA template strand with the RNAP active center (Vvedenskaya et al., 2014). The consensus sequence element for transcription-elongation pausing also comprises a less-highly-conserved upstream segment, G_P-8_ (Larson et al., 2014; Vvedenskaya et al., 2014; Imashimizu et al., 2015). In the paused TEC, the G_P-8_:C_P-8_ RNA:DNA base pair is located at the upstream end of the RNA-DNA hybrid and must be broken for RNAP translocation (Figure 6B; Vvedenskaya et al., 2014); sequence at these positions affects RNAP-translocation behavior through effects on RNA-DNA duplex thermal stability (Vvedenskaya et al., 2014).

The consensus sequence for initial-transcription pausing defined in this work: T_P-1_N_P_<u>Y</u>_A_G_A+1_ (Figure 6A, top), exhibits a striking resemblance to the downstream, most-highly-conserved portion of the consensus sequence for transcription-elongation pausing: <u>Y</u>_A_G_A+1_ (Figure 6A, bottom). This resemblance extends both to the sequence and the position of the RNAP-active-center A-site relative to sequence. Consistent with the conclusion that the consensus sequence for initial-transcription pausing: T_P-1_N_P_<u>Y</u>_A_G_A+1_, may be functionally related to the downstream, most-highly-conserved portion of the consensus sequence for transcription-elongation pausing, <u>Y</u>_A_G_A+1_, the consensus sequence for initial-transcription pausing contains T at position P_-2_ (Figure 6A) and two groups defining the consensus sequence for transcription-elongation pausing identified T as the optimal nucleotide at the corresponding position (Larson et al., 2014; Imashimizu et al., 2015). The observation that the consensus sequence for initial-transcription pausing does not contain a sequence corresponding to the upstream, less-highly-conserved portion of the consensus sequence for elongation, G_P-8_, is explained by the fact that the analyzed ITCs contain short RNA products and short RNA:DNA hybrids that do not extend to this upstream position.

We suggest that the resemblance of the consensus sequence for initial-transcription pausing and the downstream portion of the consensus sequence for transcription-elongation pausing is real and reflects a functional relationship. We infer that initial-transcription pausing and transcription-elongation pausing share mechanistic commonalities. Consistent with this proposal, substitution of the RNAP β-subunit residue 446 (β^D446A^), increases sequence-dependent pausing in both initial transcription (Figure 3A; Dulin et al., 2018) and transcription elongation (Vvedenskaya et al., 2014).

### Prospect

XACT-seq (<u>cross</u>link-between-<u>a</u>ctive-<u>c</u>enter-and-template sequencing) provides a high-throughput, direct, single-nucleotide-resolution readout of RNAP-active-center A-site position relative to DNA during transcription initiation and transcription elongation *in vivo*, in living cells. In this work, using the reagent that underpins XACT-seq, an RNAP derivative which, upon photo-activation *in vitro* or *in vivo*, forms covalent crosslinks with DNA at a position exactly 5-nt downstream of the RNAP-active-center A-site we: (i) defined the RNAP-active-center A-site position at p*lac*CONS (Figure 3), (ii) demonstrated that initial-transcription pausing occurs at a substantial fraction of promoter initial-transcribed region sequences *in vivo* (Figures 4, S2), (iii) showed that initial-transcription pausing can occur at each promoter position in the promoter-initial-transcribed region from +3 to +9 (Figures 4, S2), (iv) and showed that the σ finger contributes quantitatively to pausing at promoter positions +5 and +6, presumably through collision with the RNA 5ʹ end (Figures 3, 5E,F). Next, using XACT-seq and sampling a library of 4^11^ (∼4,000,000) promoter initial-transcribed-region sequences, we: (i) confirmed that initial-transcription pausing occurs *in vivo* (Figures 5, S4; Table S1), (ii) confirmed and quantified that initial-transcription pausing occurs at a substantial fraction (at least ∼20%) of initial-transcribed-region sequences (Figure 5C; Table S1), (iii) confirmed and quantified that initial-transcription pausing can occur at each position in the initial-transcribed-region sequence (high prevalence for positions +3 to +7; lower prevalence for positions +8 and +9; Figure 5C; Table S1), (iv) showed that initial-transcription pausing is determined primarily by promoter sequence (Figures 5, S4), (v) defined the consensus sequence for initial-transcription pausing as T_P-1_N_P_<u>Y</u>_A_G_A+1_ (Figure 5A), and (vi) showed that the consensus sequence for initial-transcription pausing resembles the consensus sequence for transcription-elongation pausing, suggesting that initial-transcription pausing and transcription-elongation pausing share mechanistic commonalities (Figure 6). A major finding is that initial-transcription pausing and transcription-elongation pausing appear to be fundamentally similar, with the key mechanistic difference, a “scrunching” mechanism of RNAP-active-center translocation in the former vs. a “stepping” mechanism of RNAP-active-center translocation in the latter, having no detectable effect, and the key structural difference between an ITC and a TEC, namely, the presence of the initiation factor σ in the former but not the latter, having only a quantitative, modulatory effect on initial-transcription pausing.

Here we show XACT-seq overcomes the limitations of methods that infer the RNAP-active-center A-site position from the identity of the RNA 3’ end, a process that, as described above, is subject to uncertainties due to translocation equilibration of TECs among pre-translocated, post-translocated, reverse-translocated, and hyper-translocated states. We further show XACT-seq overcomes the limitations of methods such as ChIP-seq and ChIP-exo that can report RNAP position, but only with low resolution, and which define the overall RNAP position but not the RNAP-active-center A-site position.

We applied XACT-seq to analysis of a library of 4^11^ (∼4,000,000) initial-transcribed region sequences. In this report, we focused on results for initial-transcription pausing. However, the same experiments also provide information for pausing in subsequent stages of transcription, including pausing at the moment of promoter escape and formation of a TEC, promoter-proximal σ-dependent pausing by TECs, and sequence-dependent elongation pausing by TECs. The findings for these subsequent stages of transcription will be reported elsewhere.

The approach that was applied here to a library of promoter sequences can be applied to genome-wide analysis of RNAP-active-center A-site positions *ex vivo* on isolated DNA or chromatin, or *in vivo*, in living cells, providing a basis for “RNAP profiling” analogous to existing methods for ribosome profiling (Ingolia et al., 2009). Combining XACT-seq and NET-seq for genome-wide analysis of RNAP-active-center A-site positions and RNA 3ʹ end positions, respectively, each with single-nucleotide resolution, should provide an unprecedently rich description of the sequence-dependent, factor-dependent transcriptional landscape.

## Acknowledgements

Work was supported by NIH grants GM124976 (PS), GM041376 (RHE), and GM118059 (BEN). JTW was supported by a fellowship from the Helen Hay Whitney Foundation.

## Author Contributions

Conceptualization, JTW, RHE, BEN Methodology, JTW, CP, YZ, IOV, PS Formal Analysis, JTW, CP, PS, YZ, DT Investigation, JTW, CP, IOV Writing, JTW, RHE, BEN Visualization, JTW, RHE, BEN Supervision, DT, RHE, BEN Project Administration, RHE, BEN Funding Acquisition, PS, RHE, BEN

## METHODS

### Plasmids

To generate plasmid pIA900-βD446A RNAP-β′^R1148Bpa^, we performed PCR using ∼20 ng of pIA900-RNAP-β′^R1148Bpa^ (Winkelman et al., 2015) in 12.5 µl reactions containing 0.8 µM oligo JW44 and 1 X Phusion HF Master Mix (Thermo Fischer Scientific) (95°C for 2 min; 95°C for 15 s, 55°C for 15s, 72°C for 5 min; 30 cycles; 72°C for 6 min). Next, to remove the wild-plasmid template, 20 units of DpnI (New England Biolabs) was added, the reactions was incubated at 37°C for 16 h,1 µl of the reaction was introduced into electrocompetent DH10B cells (Thermo Fischer Scientific), and cells plated on LB agar plates containing 100 µg/ml carbenicillin to select for transformants.

To generate plasmid pσ^70^-His pσ^70^Δ finger, we performed PCR using 4 ng of plasmid pσ^70^-His (Marr and Roberts, 1997) in 50 µl reactions containing 0.8 µM oligo JW268, 0.8 µM oligo JW269, and 1 X Phusion HF Master Mix (2 min at 95°C; 95°C for 15 s, 55°C for 15s, 72°C for 4 min (30 x); 2 min at 72°C). 30 units of DpnI was added, the reaction was incubated at 37°C for 16 h, and DNA was recovered using a PCR purification kit (Qiagen). The recovered DNA was treated with 10 units of T4 polynucleotide kinase (PNK) (New England Biolabs) in 1 X T4 PNK buffer containing 20 µM ATP for 30 min at 37°C, 400 units of T4 DNA ligase (New England Biolabs) was added, reactions were incubated at 16°C for 16 h, 1 µl of the reaction was introduced into electrocompetent DH10B cells, and cells plated on LB agar plates containing 50 µg/ml kanamycin, and recombinant plasmid DNA was isolated from individual transformants.

Plasmid pCDF-CP (Yu et al., 2017) contains a CloDF13 replication origin, a selectable marker conferring resistance to spectinomycin, and two BglI recognition sites that are used to introduce DNA fragments upstream of transcription terminator tR2. Plasmid pCDF-*lac*CONS (Yu et al., 2017) is a derivative of pCDF-CP containing sequences from positions -88 to +70 of p*lac*CONS inserted into BglI-digested pCDF-CP.

To generate plasmids pCDF-*lac*CONS-p5, -p6, -p7, -p8, and -p9, which contain the sequences between positions +3 and +13 shown in Figure 5D, we performed PCR using 1 ng of pCDF-*lac*CONS in 25 µl reactions containing 1 X Phusion HF Master Mix, 0.8 µM oligo JW80 and 0.8 µM oligo JW433, JW408, JW409, JW580, or JW575, respectively. 1 µl of this reaction was used as a template in PCR reactions (50 µl) containing 1 X Phusion HF Master Mix, 0.8 µM JW79, and 0.8 µM JW80. Amplicons were treated with BglI and BglI-digested fragments were ligated into BglI-digested pCDF-CP. The ligation mixture was transformed into NEB 5-alpha competent cells (New England Biolabs), cells plated on LB agar plates containing 50 µg/ml spectinomycin, and recombinant plasmid DNA was isolated from individual transformants.

Plasmid-borne promoter libraries p*lac*CONS-N11 and p*lac*CONS-N20 were generated using a procedure described in (Vvedenskaya et al., 2015; Vvedenskaya et al., 2018b) that provides a “self-assembling barcode,” in which for each DNA molecule in the library a first randomized sequence in a region of interest is associated with a known corresponding second randomized sequence that serves as a barcode. The procedure involves synthesis of three oligos for use in PCR. One oligo, which serves as the template for PCR amplification, contains a first randomized sequence spanning the region of interest and a second randomized sequence that serves as a barcode. The other oligos serve as amplification primers. One of these oligos contains 5’-end sequences that introduce a BglI recognition sequence and the other oligo contains 3’-end sequences complementary to the template oligo. Libraries p*lac*CONS-N11, containing up to 4^11^ initial-transcribed region sequences from position +3 to position +13, was constructed using oligo JW153 as the template and oligos s1219 and s1220 as amplification primers. Library p*lac*CONS-N20, containing up to 4^20^ initial-transcribed region sequences from position +2 to position +21, was constructed using oligo JW203 as the template and oligos s1219 and s1220 as amplification primers. Amplicons were treated with BglI and BglI-digested fragments were ligated into BglI-digested pCDF-CP. The ligation mixture was transformed into NEB 5-alpha competent cells, cells plated on LB agar plates containing 50 µg/ml spectinomycin, and recombinant plasmid DNA was isolated from ∼10^7^ transformants.

Plasmid pEVOL-pBpF contains genes directing the synthesis of an engineered Bpa-specific UAG-suppressor tRNA and an engineered Bpa-specific aminoacyl-tRNA synthetase that charges the amber suppressor tRNA with Bpa (Addgene; Chin et al., 2002).

### Proteins

Bpa-containing RNAP core enzyme derivatives RNAP-β′^R1148Bpa^ and βD446A RNAP-β′^R1148Bpa^ were prepared from *E. coli* strain NiCo21(DE3) (New England Biolabs) containing plasmid pEVOL-pBpF and plasmid pIA900-RNAP-β′^R1148Bpa^ (Winkelman et al., 2015) or plasmid pIA900-βD446A RNAP-β′^R1148Bpa^, using procedures described in Winkelman et al., 2015.

Wild-type σ^70^ or a σ^70^ derivative containing a deletion of the σ finger (residues 513-519), were prepared from *E. coli* strain NiCo21(DE3) containing plasmid pσ^70^-His (gift of J. Roberts; Marr and Roberts, 1997) or plasmid pσ^70^-His Δ finger using procedures described in (Marr and Roberts, 1997). To form RNAP holoenzyme, 1 μM RNAP core enzyme and 5 μM wild-type σ^70^ or Δ finger σ^70^ in 10 mM Tris-Cl (pH 8.0), 100 mM KCl, 10 mM MgCl_2_, 0.1 mM EDTA, 1 mM DTT, and 50% glycerol were incubated for 30 min at 25°C.

### Oligonucleotides

Oligodeoxyribonucleotides (Table S2) were purchased from IDT. Diribonucleotide ApA (HPLC-purified) was purchased from TriLink Biotechnologies.

### Templates for *in vitro* assays

Linear DNA templates used for *in vitro* transcription assays contain sequences from positions -88 to +70 of p*lac*CONS (Figure 3A,B) or positions -88 to +70 of p*lac*CONS-p5, p*lac*CONS-p6, p*lac*CONS-p7, p*lac*CONS-p8, or p*lac*CONS-p9 (Figure 5D-F). Templates were generated by PCR in reactions containing 1 X Phusion HF Master Mix, 0.8 µM primer JW61, 0.8 µM primer JW62, and ∼1 pg of plasmids p*lac*CONS-p5, -p6, -p7, -p8, or -p9. Reaction products were purified using a PCR purification kit.

### Determination of RNAP-active-center A-site positions by protein-DNA photo-crosslinking *in vitro*

*In vitro* photo-crosslinking and crosslink mapping experiments were done using procedures described in (Yu et al., 2017). For the experiments in Figure 3A,B, 50 μl reactions containing 20 nM RNAP holoenzyme, 4 nM template, 100 µM ApA dinucleotide (where indicated), and 1 X RB [10 mM Tris-Cl, pH 8.0, 70 mM NaCl, 10 mM MgCl_2_, and 0.1 mg/ml bovine serum albumin (BSA)] were incubated for 5 min at 25°C, 5 uL of 100 µM ATP, 100 µM CTP, 100 µM GTP, and 100 µM UTP (final concentration of 9 µM ATP, 9 µM CTP, 9 µM GTP, and 9 µM UTP) were added (where indicated), reactions were incubated 2 min at 25°C, and subjected to UV irradiation for 2 min at 25°C in a Rayonet RPR-100 photochemical reactor equipped with 16 x 350 nm tubes (Southern New England Ultraviolet). For the experiments in Figure 5C, 50 μl reactions containing 20 nM RNAP holoenzyme, 4 nM template, and 1 X RB were incubated for 5 min at 25°C, 5 uL of 1 mM ATP, 1 mM CTP, 1 mM GTP, and 1 mM UTP were added, reactions were incubated 2 min at 25°C, and subjected to UV irradiation for 2 min at 25°C. For the experiments in Figure 5D, 50 μl reactions containing 50 nM RNAP holoenzyme, 20 nM template (4 nM of p*lac*CONS-p5, -p6, -p7, -p8, and -p9), and 1 X RB were incubated for 5 min at 25°C, 5 μL of 1 mM ATP, 1 mM CTP, 1 mM GTP, and 1 mM UTP was added, reactions were incubated 2 min at 25°C, and subjected to UV irradiation for 2 min at 25°C.

To denature RNAP-DNA complexes, reactions were mixed with 15 μl 5 M NaCl and 6 μl 100 µg/ml heparin, incubated for 5 min at 95°C and then cooled to 4°C. Crosslinked RNAP-DNA complexes were isolated by adding 20 µl MagneHis Ni-particles (Promega) equilibrated and suspended in 10 mM Tris-Cl, pH 8.0, 1.2 M NaCl, 10 mM MgCl_2_, 10 μg/ml heparin, and 0.1 mg/ml BSA; MagneHis Ni-particles were collected using a magnetic microfuge tube rack; particles were washed with 50 µl 10 mM Tris-Cl, pH 8.0, 1.2 M NaCl, 10 mM MgCl_2_, 10 μg/ml heparin, and 0.1 mg/ml BSA, washed twice with 50 µl 1 X *Taq* DNA polymerase buffer (New England Biolabs), and particles (which contained bound RNAP-DNA complexes) were resuspended in 10 µl 1 X *Taq* DNA polymerase buffer. Primer extension reactions (12.5 µl) were performed by combining 2 µl of the recovered RNAP-DNA complexes, 1 µl of 1 µM ^32^P-5’-end-labeled primer JW61 [200 Bq/fmol; prepared using [γ^32^P]-ATP (PerkinElmer) and T4 polynucleotide kinase (New England Biolabs) as described in (Sambrook et al., 2006)], 1 μl 10 X dNTPs (2.5 mM dATP, 2.5 mM dCTP, 2.5 mM dGTP, 2.5 mM TTP, 0.5 μl 5 U/μl Taq DNA polymerase (New England Biolabs), 5 μl 5 M betaine, 0.625 μl 100% dimethyl sulfoxide, and 1.25 µl 10 X *Taq* DNA polymerase buffer; 40 cycles of 30 s at 95°C, 30 s at 53°C, and 30 s at 72°C. Reactions were stopped by addition of 12.5 μl 1 X TBE, 8 M urea, 0.025% xylene cyanol, and 0.025% bromophenol blue; radiolabeled products were separated by electrophoresis on 10% 8M urea slab gels (equilibrated and run in 1 X TBE) and visualized by storage-phosphor imaging (Typhoon 9400 variable-mode imager; GE Life Science). Positions of RNAP-DNA crosslinking were determined by comparison to products of a DNA-nucleotide sequencing reaction generated using oligo JW61 and a linear DNA template containing sequences from positions -88 to +70 of p*lac*CONS (Thermo Sequenase Cycle Sequencing Kit; Affymetrix).

### Determination of RNAP-active-center A-site positions by protein-DNA photo-crosslinking *in vivo*

*In vivo* photo-crosslinking and crosslink mapping experiments of Figures 3B, 4B, and S5 were done essentially as described in (Yu et al., 2017) except the RNAP active site was not mutationally inactivated. Analysis of RNAP-active-center A-site positions *in vivo* for p*lac*CONS transcription complexes was performed by sequential introduction of plasmid pCDF-*lac*CONS, plasmid pIA900-RNAP-β′^R1148Bpa^, and plasmid pEVOL-pBpF into *E. coli* strain NiCo21(DE3) by transformation. After the final transformation step, cells were plated on LB agar containing 100 μg/ml carbenicillin, 50 μg/ml spectinomycin, 50 μg/ml streptomycin, and 25 μg/ml chloramphenicol; at least 1,000 individual colonies were scraped from the plate, combined, and used to inoculate 250 ml LB broth containing 1 mM Bpa (Bachem), 100 μg/ml carbenicillin, 50 μg/ml spectinomycin, 50 μg/ml streptomycin, and 25 μg/ml chloramphenicol in a 1000 ml flask (Bellco) to yield OD_600_ = 0.3; the culture was placed in the dark and shaken (220 rpm) for 1 h at 37°C; isopropyl-β-D-thiogalactoside (IPTG) was added to 1 mM; and the culture was placed in the dark and shaken (220 rpm) for 3 h at 37°C.

Analysis of RNAP-active-center A-site positions *in vivo* for p*lac*CONS-N11 and p*lac*CONS-N20 transcription complexes was performed by sequential introduction into *E. coli* strain NiCo21(DE3) of the pCDF-*lac*CONS-N11 library (yielding ∼2 million transformants) or the p*lac*CONS-N20 library (yielding ∼2 million transformants), plasmid pIA900-RNAP-β′^R1148Bpa^ (yielding ∼15 million transformants), and plasmid pEVOL-pBpF (yielding ∼4 million transformants). After the final transformation step, cells were plated on ∼10-15 LB agar plates containing 100 μg/ml carbenicillin, 50 μg/ml spectinomycin, 50 μg/ml streptomycin, and 25 μg/ml chloramphenicol to yield a lawn. Colonies were scraped from the surface of the plates, combined, and used to inoculate 250 ml LB broth containing 1 mM Bpa, 100 μg/ml carbenicillin, 50 μg/ml spectinomycin, 50 μg/ml streptomycin, and 25 μg/ml chloramphenicol in a 1000 ml flask to yield OD_600_ = 0.3; the culture was placed in the dark and shaken (220 rpm) for 1 h at 37°C; IPTG was added to 1 mM; and the culture was placed in the dark and shaken (220 rpm) for 3 h at 37°C.

To enable analysis of static transcription complexes, a portion of the cell cultures containing pCDF-*lac*CONS, pCDF-*lac*CONS-N11, or pCDF-*lac*CONS-N20 were removed, Rifampin was added to a final concentration of 200 µg/ml, and the culture was shaken for 20 min prior to harvesting cells for UV irradiation. Cell suspensions (7 ml) were removed from each culture to a 13 mm x 100 mm borosilicate glass test tube (VWR) and subjected to UV irradiation for 10 min at 25°C. Cells were collected by centrifugation (3000 x g; 15 min at 4°C) and cell pellets were stored at -20°C. To lyse cells, the frozen cell pellets were incubated at 4°C for 30 min, re-suspended in 40 ml of 50 mM Na_2_HPO_4_ (pH 8.0), 1.4 M NaCl, 20 mM imidazole, 14 mM β-mercaptoethanol, 0.1% Tween 20, and 5% ethanol containing 2 mg egg white lysozyme, and sonicated for 5 min at 4°C. Lysates were centrifuged (23,000 x g; 40 min at 4°C), supernatants were added to 1 ml Ni-NTA-agarose (Qiagen) in 50 mM Na_2_HPO_4_ (pH 8.0), 1.4 M NaCl, 20 mM imidazole, 0.1% Tween 20, 5 mM β-mercaptoethanol, and 5% ethanol, and the mixture was incubated 30 min at 4°C with gentle rocking. The slurry was loaded into a 15 ml polyprep column (BioRad) to collect the Ni-NTA-agarose resin. The resin was washed with 10 ml of 1 X WB (50 mM Na_2_HPO_4_, pH 8.0, 300 mM NaCl, 20 mM imidazole, 0.1% Tween 20, 5 mM β-mercaptoethanol, and 5% ethanol) and His-tagged RNAP was eluted from the resin with 3 ml of 1 X WB containing 300 mM imidazole. The eluate was concentrated to 0.2 ml in 20 mM Tris-Cl (pH 8.0), 200 mM KCl, 20 mM MgCl_2_, 0.2 mM EDTA, and 1 mM DTT using a 1000 MWCO Amicon Ultra-4 centrifugal filter (EMD Millipore), 0.2 ml glycerol was added, and the sample was stored at -20°C.

To denature RNAP-DNA complexes, 25 μl of the concentrated eluate was mixed with 25 μl water, 15 μl 5 M NaCl, and 6 μl 100 µg/ml heparin, incubated at 95°C for 5 min, then incubated at 4°C for 30 min. Crosslinked RNAP-DNA complexes were isolated by adding 20 µl MagneHis Ni-particles equilibrated and suspended in 10 mM Tris-Cl, pH 8.0, 1.2 M NaCl, 10 mM MgCl_2_, 10 μg/ml heparin, and 0.1 mg/ml BSA; MagneHis Ni-particles were collected using a magnetic microfuge tube rack; particles were washed with 50 µl 10 mM Tris-Cl, pH 8.0, 1.2 M NaCl, 10 mM MgCl_2_, 10 μg/ml heparin, and 0.1 mg/ml BSA, washed twice with 50 µl 1 X *Taq* DNA polymerase buffer (New England Biolabs), and the particles (which contained bound RNAP-DNA complexes) were resuspended in 10 µl 1 X *Taq* DNA polymerase buffer. Primer extension reactions and analysis of radiolabeled products generated in these reactions were performed using procedures identical to those used in the analysis of p*lac*CONS *in vitro* (see above) using material isolated from cells containing pCDF-*lac*CONS (Figure 3B), cells containing the pCDF-*lac*CONS-N11 library (Figure 4B), or cells containing the pCDF-*lac*CONS-N20 library (Figure S2).

XACT-seq experiments (see below) were performed using denatured RNAP-DNA complexes isolated from cells containing the pCDF-*lac*CONS-N11 library.

### XACT-seq: primer extension

Primer extension reactions (50 µl) were performed by combining 8 µl of recovered RNAP-DNA complexes, 1 µl of 1 µM primer s128a, 5 μl 10 X dNTPs (2.5 mM dATP, 2.5 mM dCTP, 2.5 mM dGTP, 2.5 mM TTP, 1 μl 5 U/μl Taq DNA polymerase, 10 μl 5 M betaine, 5 μl 100% dimethyl sulfoxide, and 5 µl 10 X Taq DNA polymerase buffer, and cycling 40 times through 30 s at 95°C, 30 s at 53°C, and 30 s at 72°C. Primer extension products were isolated by ethanol precipitation, washed twice with 80% cold ethanol, resuspended in 20 µl water, and mixed with 20 µl of 2 X RNA loading dye (95% deionized formamide, 18 mM EDTA, 0.25% SDS, xylene cyanol, bromophenol blue, amaranth).

Primer extension products were separated by electrophoresis on 10% 7M urea slab gels (equilibrated and run in 1 X TBE). The gel was stained with SYBR Gold nucleic acid gel stain (Life Technologies) and ssDNA products ∼35- to ∼75-nt in size were excised from the gel. To elute nucleic acids from the gel, the gel fragment was crushed as described in (Vvedenskaya et al., 2018a), 300 μl of 0.3 M NaCl in 1 X TE buffer was added, the mixture was incubated for 10 min at 70°C, and the supernatant was collected using a Spin-X column (Corning). The elution procedure was repeated, supernatants were combined, and nucleic acids were recovered by ethanol precipitation, washed twice with 80% cold ethanol, and resuspended in 5 μl of nuclease-free water.

### XACT-seq: 3′-adapter ligation and library amplification

The recovered primer extension products (5 μl) were combined with 1 μl 10 X NEB buffer 1, ∼0.8 μM 3′-adapter oligo s1248 [5′ adenylated and 3′-end blocked oligo containing ten randomized nucleotides (10N) at the 5′ end], 5 mM MnCl_2_ and 1 μM of 5′-AppDNA/RNA ligase (New England Biolabs) in a final volume of 10 μl. The mixture was incubated for 1 h at 65°C followed by 3 min at 90°C, and cooled to 4°C for 5 min. The reaction was combined with 15 μl of mixture containing 10 U of T4 RNA ligase 1 (New England Biolabs), 1 X T4 RNA ligase 1 reaction buffer, 12% PEG 8000, 10 mM DTT, 60 μg/mL BSA. Reactions were incubated at 16°C for 16 h.

To determine the effect of sequence on the efficiency of ligation between primer extension products and the 3′-adapter oligo s1248, we performed a control ligation reaction containing oligo s1248 and oligo JW402, which contains 15 random nucleotides (15N) at the 5ʹ end.

Adapter-ligated products were separated by electrophoresis on 10% 7M urea slab gels (equilibrated and run in 1 X TBE). The gel was stained with SYBR Gold nucleic acid gel stain and species ranging from ∼60 to ∼100-nt (for reactions containing oligo s1248 and primer extension products) or ∼50 and 90 nt (for reactions containing oligo s1248 and oligo JW402) were isolated by gel excision. The gel fragment was crushed, 600 μl of 0.3M NaCl in 1 X TE buffer was added, the mixture was incubated for 2 h at 37°C, the supernatant was collected using a Spin-X column (Corning). The elution procedure was repeated, supernatants were combined, and nucleic acids were recovered by ethanol precipitation, washed twice with 80% cold ethanol, and resuspended in 20 μl of nuclease-free water.

Adapter-ligated DNA (5 μl) was added to PCR reactions containing 1 X Phusion HF reaction buffer, 0.2 mM dNTPs, 0.25 μM Illumina RP1 primer, 0.25 μM Illumina RPI index primer and 0.02 U/μl Phusion HF polymerase [40 μl total volume; 98°C for 30 s, 98°C for 10 s, 62°C for 20 s, 72°C for 10 seconds (11 cycles), 72°C for 5 min]. Reaction products were separated by electrophoresis on a non-denaturing 10% slab gel (equilibrated and run in 1 X TBE), and amplicons between ∼160 bp and ∼170 bp were isolated by gel excision. The gel fragment was crushed, 600 μl of 0.3M NaCl in 1 X TE buffer was added, the mixture was incubated for 2 h at 37°C, the supernatant was collected using a Spin-X column. The elution procedure was repeated, supernatants were combined, and nucleic acids were recovered by ethanol precipitation, washed twice with 80% cold ethanol, and resuspended in 20 μl of nuclease-free water.

Libraries generated by this procedure are: CP21/CP21D, CP23/CP23D, CP27/CP27C, CP22/CP22D, CP24/CP24D, and CP28/CP28C.

### XACT-seq: generation of library for analysis of template sequences in the p*lac*CONS-N11 library

We used emulsion PCR (ePCR) to generate a DNA library to identify template sequences present in the p*lac*CONS-N11 library. ePCR reaction mixtures contained ∼10^9^ molecules of the p*lac*CONS-N11 plasmid library, 1 X Detergent-free Phusion HF reaction buffer containing 5 μg/ml BSA, 0.4 mM dNTPs, 0.5 μM Illumina RP1 primer, 0.5 μM Illumina index primer and 0.04 U/μl Phusion HF polymerase [95°C for 10 s, 95°C for 5 s, 60°C for 5 s, 72°C for 15 seconds (20 cycles), 72°C for 5 min]. DNA was isolated from ePCR reactions using a Micellula DNA Emulsion and Purification Kit. The emulsion was broken, DNA was purified according to the manufacturer’s recommendations, recovered by ethanol precipitation, and resuspended in 20 μl of nuclease-free water. Products were subjected to electrophoresis on a non-denaturing 10% slab gel (equilibrated and run in 1 X TBE), the 204 bp fragment was excised from the gel. The gel fragment was crushed, 600 μl of 0.3M NaCl in 1 X TE buffer was added, the mixture was incubated for 2 h at 37°C, the supernatant was collected using a Spin-X column. The elution procedure was repeated, supernatants were combined, and nucleic acids were recovered by ethanol precipitation, washed twice with 80% cold ethanol, and resuspended in 20 μl of nuclease-free water. The library generated by this procedure is CP26T.

### XACT-seq: quantitation of template sequences

To determine the relative abundance of each template sequences, we isolated plasmid DNA from 2 ml of cell suspensions harvested prior to UV irradiation. Plasmid DNA (∼120 ng) was mixed with 1 X Phusion HF reaction buffer, 0.1 mM dNTPs, 0.125 μM biotinylated oligonucleotide JW401, and 0.01 U/μl Phusion HF polymerase in a 20 μl reaction. Primer extension reactions we performed by denaturing plasmid DNA for 30 s at 98°C, followed by 2 cycles of denaturation for 10 s at 98°C, annealing for 20 s at 62°C, extension for 15 s at 72°C, and a final extension for 5 min at 72°C. Reactions were subjected to electrophoresis using a non-denaturing 10% slab gel (equilibrated and run in 1 X TBE). The gel was stained with SYBR Gold nucleic acid gel stain and the region of the gel containing ssDNA products larger than ∼40-nt was excised. The gel fragment was crushed, 600 μl of 0.3M NaCl in 1 X TE buffer was added, the mixture was incubated for 2 h at 37°C, the supernatant was collected using a Spin-X column. The elution procedure was repeated, supernatants were combined, and nucleic acids were recovered by ethanol precipitation, washed twice with 80% cold ethanol, and resuspended in 10 μl of nuclease-free water. 1 μl of this solution was amplified by ePCR, products were recovered from emulsions as described above, and subjected to electrophoresis on a non-denaturing 10% slab gel (equilibrated and run in 1 X TBE). The 204 bp fragment was excised from the gel, the gel fragment was crushed, 600 μl of 0.3M NaCl in 1 X TE buffer was added, the mixture was incubated for 2 h at 37°C, the supernatant was collected using a Spin-X column. The elution procedure was repeated, supernatants were combined, and nucleic acids were recovered by ethanol precipitation, washed twice with 80% cold ethanol, and resuspended in 20 μl of nuclease-free water.

Libraries generated by this procedure are: Vv1362, Vv1363, and Vv1364.

### XACT-seq: HTS sample serial numbers

CP21, CP21D, CP23, CP23D, CP27, and CP27C are samples used for the identification of RNAP-active-center A-site positions in active transcription complexes *in vivo* (-Rifampin). CP22, CP22D, CP24, CP24D, CP28, and CP28C are samples used for the identification of RNAP-active-center A-site positions in static transcription complexes *in vivo* (+ Rifampin). CP21/CP21D, CP23/CP23D, CP27/CP27C, CP22/CP22D, CP24/CP24D, and CP28/CP28C are technical replicates derived from the same biological sample.

Sample CP26T was used to identify template sequences present in the p*lac*CONS-N11 library. Vv1362, Vv1363, and Vv1364 are samples used quantify template sequences for each biological replicate (Vv1362 was used to quantify template sequences for CP21/CP21D/CP22/CP22D; Vv1363 was used to quantify template sequences for CP23/CP23D/ CP24/CP24D; Vv1364 was used to quantify template sequences for CP27/CP27C/CP28/CP28C). Samples CP40, CP41, and CP42 are libraries generated to determine the effect of sequence on the efficiency of adapter ligation.

All samples were sequenced using an Illumina NextSeq with custom sequencing primer s1115.

### XACT-seq: data analysis

Sequencing reads derived from sample CP26T that were a perfect match to non-randomized sequences of p*lac*CONS-N11 were identified: aggc**ttgaca**ctttatgcttcggctcg**tataat**gtgtggaa<u>nnnnnnnnnnn</u>gataacaatttcaacaat<u>nnnnnnnnnnnnnnnnn</u>tggaattctcgggtgccaagg; -35 element and -10 element are in bold; randomized sequences from +3 to +13 (“region of interest”) and from +32 to +48 (“barcode”) are underlined. From these reads, the sequences of the region of interest (+3 to +13) and barcode (+32 to +48) were used to identify template sequences containing a barcode that was associated with a unique sequence in the region of interest. These barcodes were used to assign sequencing reads derived from primer extension products (samples CP21/CP21D, CP23/CP23D, CP27/CP27C, CP22/CP22D, CP24/CP24D, and CP28/CP28C) to a p*lac*CONS-N11 template sequence.

Positions of RNAP-DNA crosslinking on each p*lac*CONS-N11 template sequence were identified and quantified from the analysis of samples CP21/CP21D, CP23/CP23D, CP27/CP27C, CP22/CP22D, CP24/CP24D, and CP28/CP28C using procedures described in (Winkelman et al., 2016b). For each read, the sequence of the barcode was used to define the p*lac*CONS-N11 template from which the primer extension product was derived and the sequence of the 3′-end of primer extension product was used to define the position of crosslinking (position X) and position of the RNAP-active-center A-site (the position 5 bp upstream of X, X-5). For each p*lac*CONS-N11 template sequence, we determined the read count (RC) for extension products derived from transcription complexes for which the RNAP-active-center A-site was at each position from -3 to +20 (RC_-3_ to RC_+20_; corresponding to RNAP-DNA crosslinking at each position from +3 to +25). The RNAP occupancy for promoter-position X over a region of initial-transcribed region sequence from promoter-position Y to promoter position Z is defined as the value of RC_X_/RC_total_, where RC_total_ represents the total read count for all A-site positions from position Y to position Z.

### XACT-seq: identification of sequence determinants for initial-transcription pausing

To define the global consensus sequence for initial-transcription pausing, we calculated the RNAP occupancy value for promoter-positions +3, +4, +5, +6, +7, +8, or +9 over sequences from position -3 to position +20 for each p*lac*CONS-N11 template sequence. Pause sites were identified as sequences having RNAP occupancy values at positions +3, +4, +5, +6, +7, +8, or +9 that were in the top 1% of all values. The global consensus sequence for initial-transcription pausing (Figure 5A) was derived by aligning nontemplate-strand sequences of these pause sites in the following manner: the sequences of nontemplate-strand nucleotides of pause sites with the RNAP-active-center A-site at positions +3, +4, +5, +6, +7, +8, or +9 were aligned to generate the sequence logo (Wagih, 2017) for the A-site position (A), the position 1 bp downstream of the A-site (A_+1_), and the position 2 bp downstream of the A-site (A_+2_); the sequences of nontemplate-strand nucleotides of pause sites with the RNAP-active-center A-site at positions +4, +5, +6, +7, +8, or +9 were aligned to generate the sequence logo for the P-site position (P); the sequences of nontemplate-strand nucleotides of pause sites with the RNAP-active-center A-site at positions +5, +6, +7, +8, or +9 were aligned to generate the sequence logo for the position immediately upstream of the P-site (P_-1_); the sequences of nontemplate-strand nucleotides of pause sites with the RNAP-active-center A-site at positions +6, +7, +8, or +9 were aligned to generate the sequence logo for the position 2 bp upstream of the P-site (P_-2_); the sequences of nontemplate-strand nucleotides of pause sites with the RNAP-active-center A-site at positions +7, +8, or +9 were aligned to generate the sequence logo for the position 3 bp upstream of the P-site (P_-3_); the sequences of nontemplate-strand nucleotides of pause sites with the RNAP-active-center A-site at positions +8, or +9 were aligned to generate the sequence logo for the position 4 bp upstream of the P-site (P_-4_); the sequences of nontemplate-strand nucleotides of pause sites with the RNAP-active-center A-site at position +9 were aligned to generate the sequence logo for the position 5 bp upstream of the P-site (P_-5_).

To generate the plot shown in Figure 5B, values of RNAP occupancy for promoter-positions +5, +6, +7, +8, or +9 over sequences from position +5 to position +9 were calculated for nontemplate-strand tetranucleotide sequences at the positions corresponding to P_-1_PAA_+1_ (+3 to +6 for A-site position +5; +4 to +7 for A-site position +6; +5 to +8 for A-site position +7; +6 to +9 for A-site position +8; +7 to +10 for A-site position +9). Plot shows the mean and SEM calculated from three biological replicates.

To define the consensus sequence for initial-transcription pausing at each position (Figure 5C), we identified p*lac*CONS-N11 template sequences for which RC_total_ >10 (with RC_total_ = sum of RC at each position from -3 to +20). Pause sites were identified as positions with RNAP occupancy >50%. The position-sequence logos for pausing at positions +3, +4, +5, +6, +7, +8, or +9 were generated by aligning the sequences of nontemplate-strand nucleotides of pause sites with the RNAP-active-center A-site at positions +3, +4, +5, +6, +7, +8, or +9, respectively.

---

**Figure S1.**
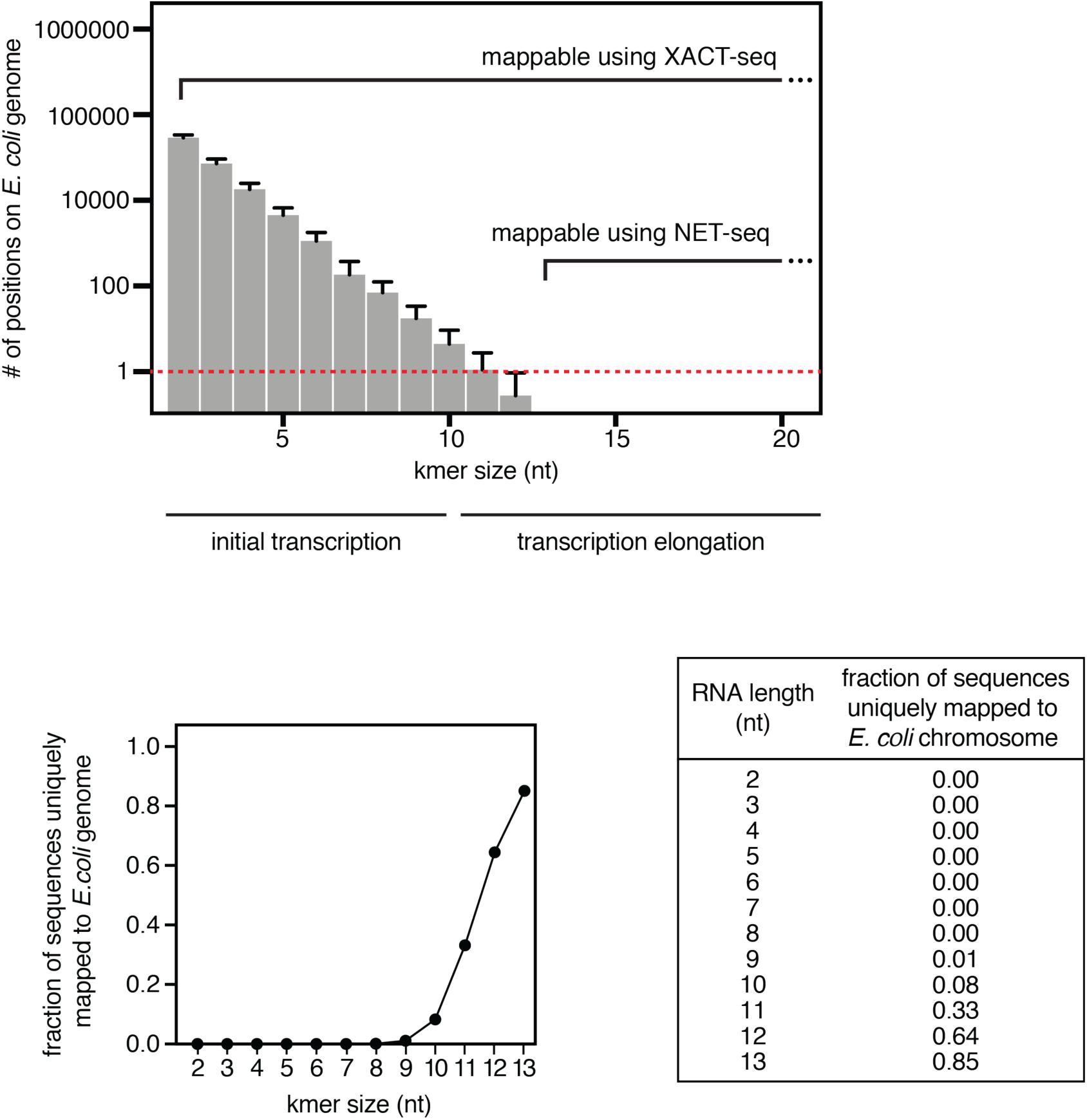
Inapplicability of RNA-based sequencing methods, such as NET-seq, for analysis of initial transcription. Relationship between RNA product length and number of occurrences of sequence on the *E. coli* genome. Top, plot of mean ± SD of the number of occurrences of sequences on the *E. coli* genome as a function of RNA length. Bottom, fraction of sequences uniquely mappable to the *E. coli* genome as a function of RNA length. Because essentially all RNA products ≤ 10 nt, and a substantial number of RNA products ≤ 13 nt, have more than one occurrence on the *E. coli* genome, and because many RNA product sequences ≤ 13 nt are not uniquely mappable to the *E. coli* genome, RNA-based sequencing methods cannot be used to study initial transcription. In contrast, because RNAP-active-center A-site positions for RNA products of any length are mappable to the *E. coli* genome using XACT-seq, this method can be used for analysis of initial transcription.

**Figure S2.**
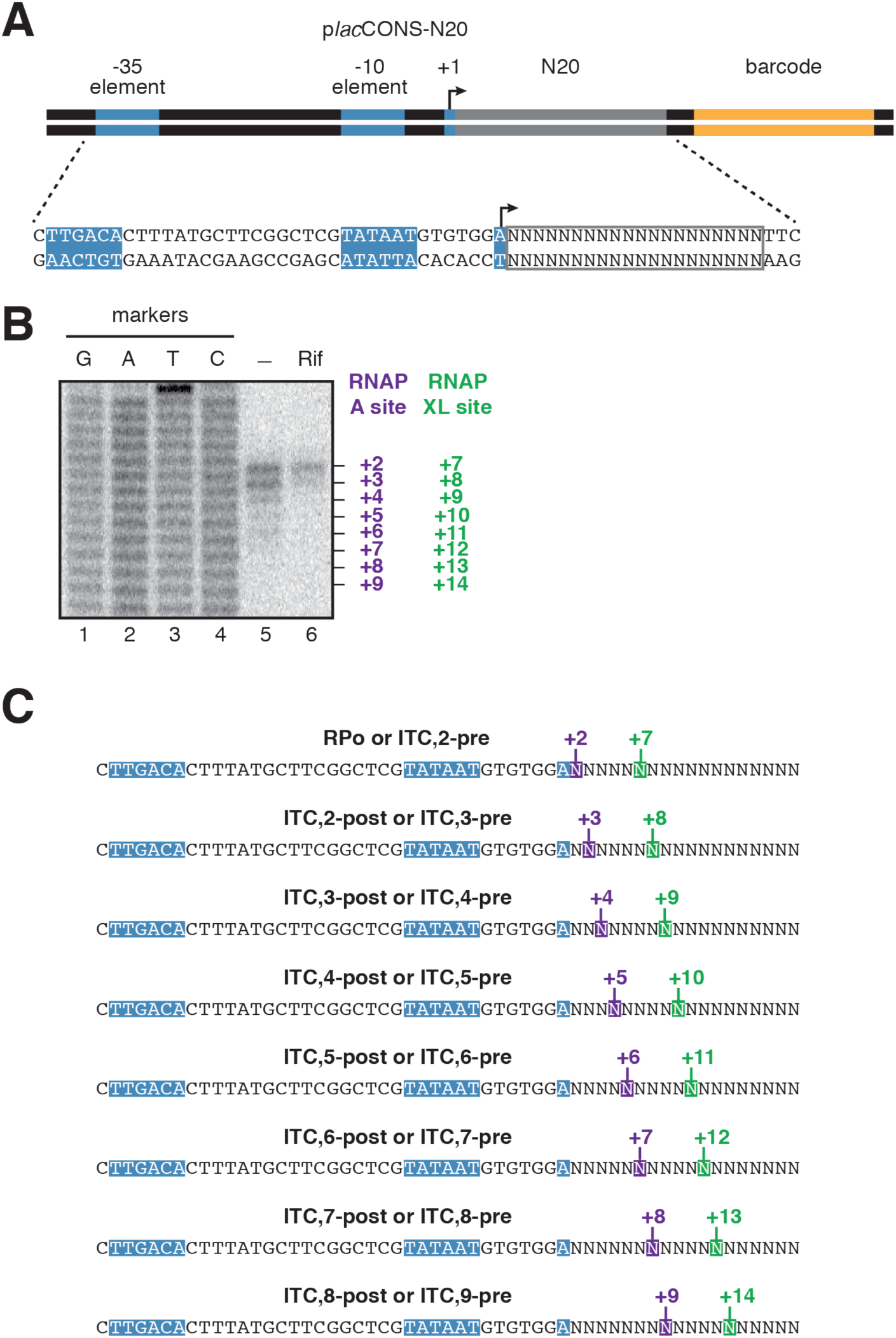
Promoter-position dependence of initial-transcription pausing for a library of up to 4^20^ promoter sequences *in vivo*. (A) p*lac*CONS template library containing all possible sequences from promoter position +2 to promoter position +21 (p*lac*CONS-N20). (B) Left, PAGE analysis of RNAP-active-center A-site positions *in vivo*. Right, histogram showing signals detected in the absence (black line) or presence (grey line) of Rifampin. Markers, DNA sequence ladder generated using p*lac*CONS-N20. (C) Position of RNAP-active-center A-site (purple) and nucleotide crosslinked to Bpa (green) defined relative to the TSS position, +1.

**Figure S3.**
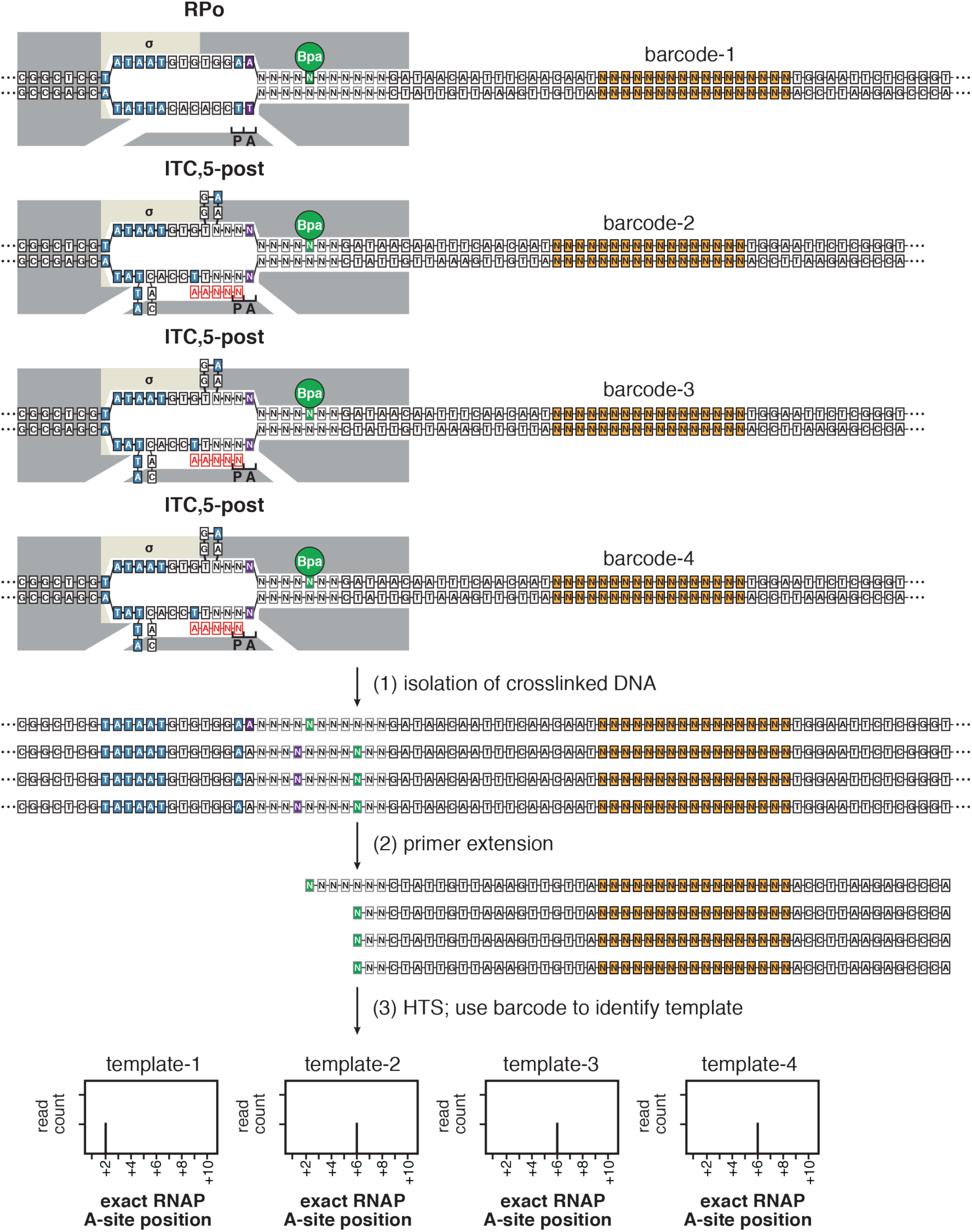
Identification of RNAP-active-center A-site positions in initial transcription by XACT-seq. Identification of RNAP-active-center A-site positions for transcription complexes formed on p*lac*CONS-N11 template sequences. Grey boxes, nucleotides of the randomized initial-transcribed region (+3 to +13); orange-filled boxes, nucleotides of the randomized barcode region (+32 to +48). Other colors and symbols as in Figure 3C.

**Figure S4.**
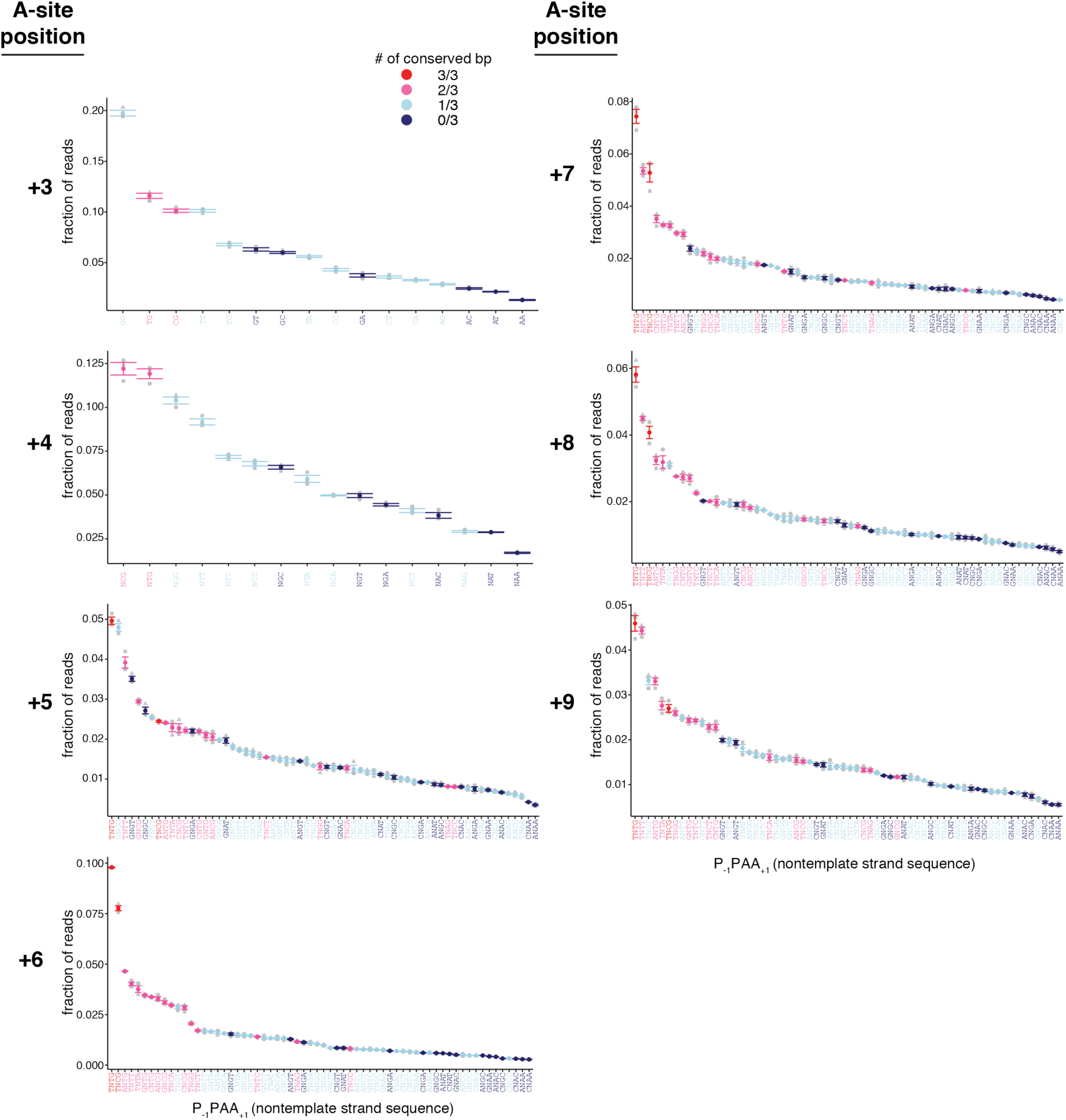
Promoter-sequence determinants for initial-transcription pausing in a library of 4^11^ (∼4,000,000) promoters *in vivo*: additional data. RNAP occupancy at each dinucleotide sequence (for RNAP-active-center A-site position +3), at each trinucleotide sequence (for RNAP-active-center A-site position +4), or at each tetranucleotide sequence (for RNAP-active-center A-site positions +5 to +9). Colors as in Figure 5C. Mean ± SEM (n = 3).

**Figure S5.**
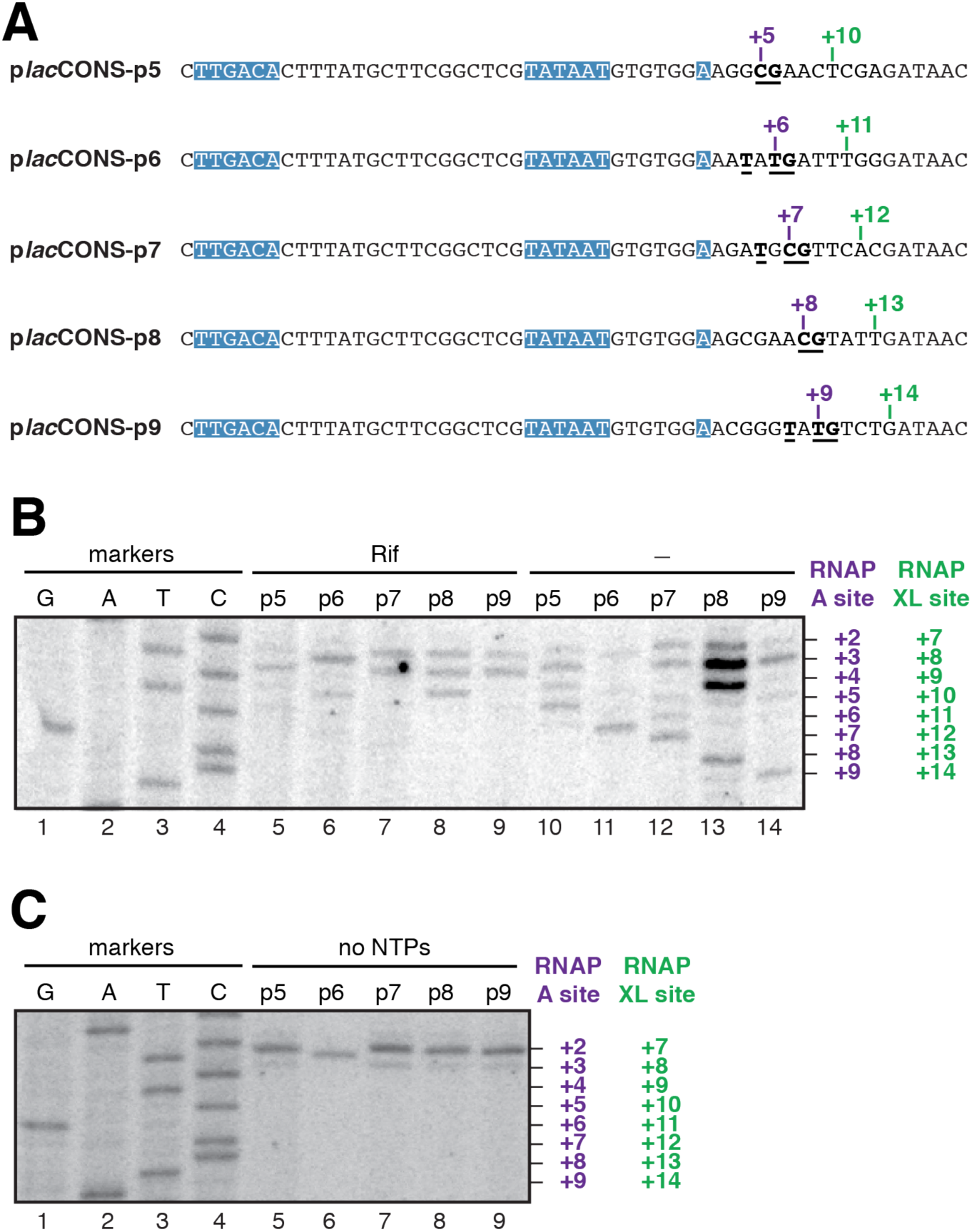
Promoter-sequence determinants for initial-transcription pausing in a library of 4^11^ (∼4,000,000) promoters *in vivo*: additional data. (A) Representative sequences yielding high RNAP occupancy at RNAP-active-center A-site positions +5, +6, +7, +8, or +9, respectively. Colors as in Figure 4C. (B) RNAP-active-center A-site positions and crosslinking positions *in vivo* for the sequences of (A) with high RNAP occupancy at RNAP-active-center A-site positions +5, +6, +7, +8, or +9. Lanes 1-4 present sequence markers, lanes 5-9 present data for wild-type RNAP in the presence of the RNAP inhibitor Rifampin, and lanes 10-14 present data for wild-type RNAP in the absence of the RNAP inhibitor Rifampin. (C) RNAP-active-center A-site positions and crosslinking positions *in vitro* for the sequences of (A) with high RNAP occupancy at RNAP-active-center A-site positions +5, +6, +7, +8, +9. Lanes 1-4 present sequence markers, lanes 5-9 present data for wild-type RNAP in the absence of NTPs.

**Table S1.**
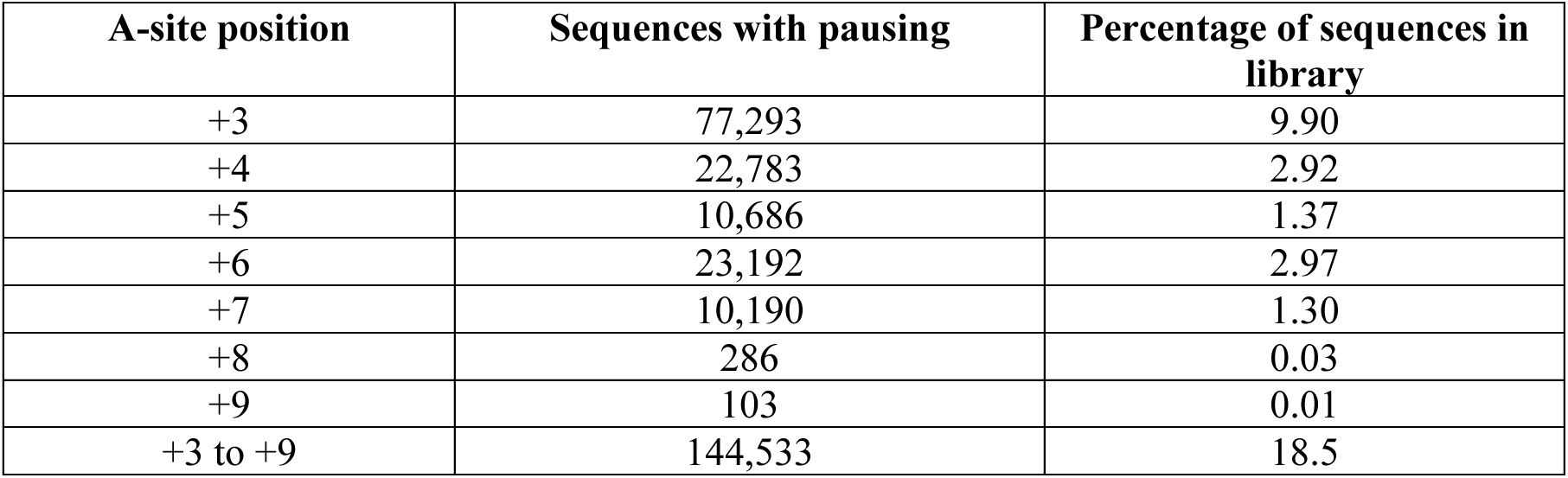
p*lac*CONS-N11 template sequences containing pauses at positions +3 to +9 identified by XACT-seq.

| A-site position | Sequences with pausing | Percentage of sequences in library |
| --- | --- | --- |
| +3 | 77,293 | 9.90 |
| +4 | 22,783 | 2.92 |
| +5 | 10,686 | 1.37 |
| +6 | 23,192 | 2.97 |
| +7 | 10,190 | 1.30 |
| +8 | 286 | 0.03 |
| +9 | 103 | 0.01 |
| +3 to +9 | 144,533 | 18.5 |

**Table S2.**
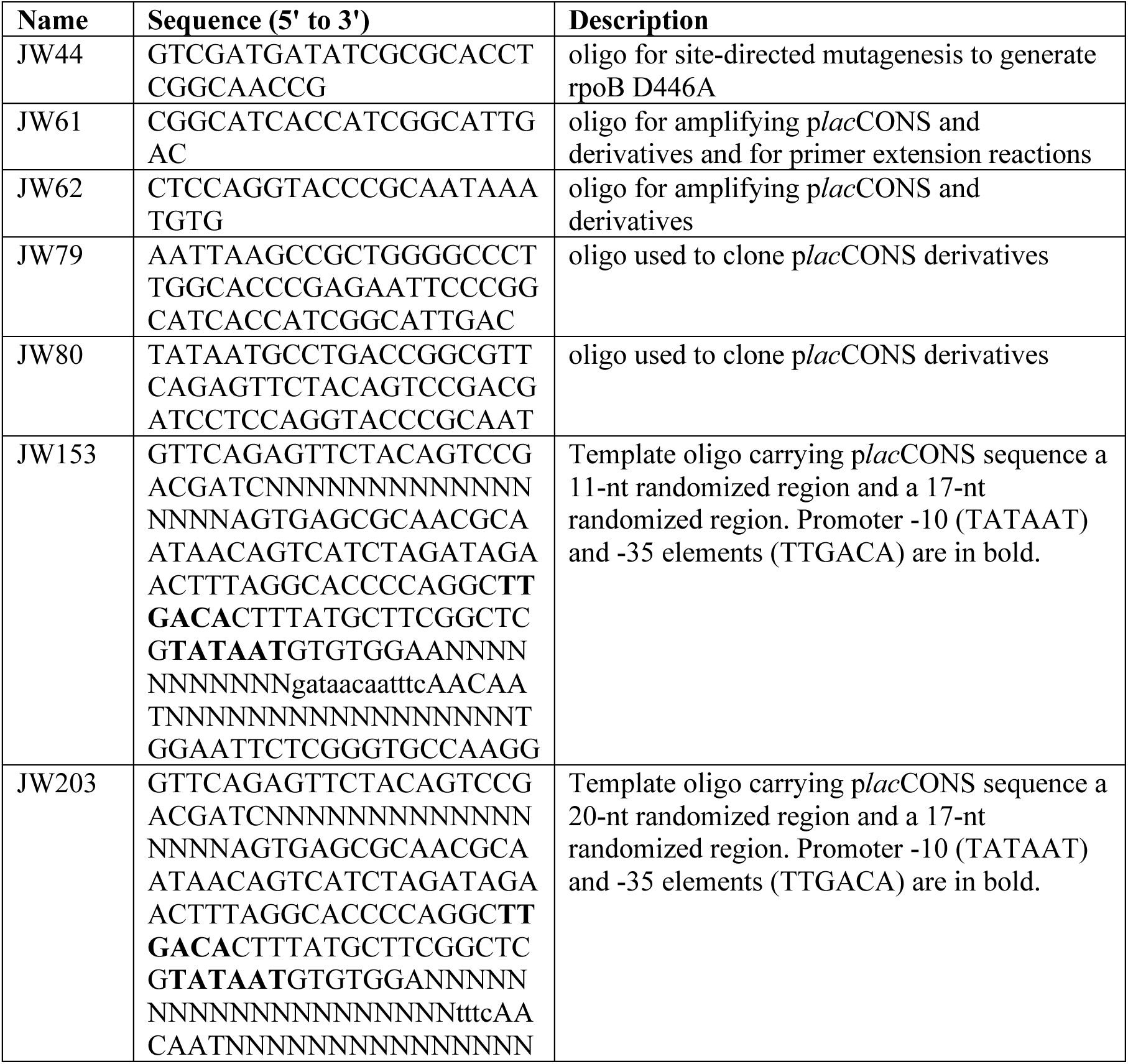
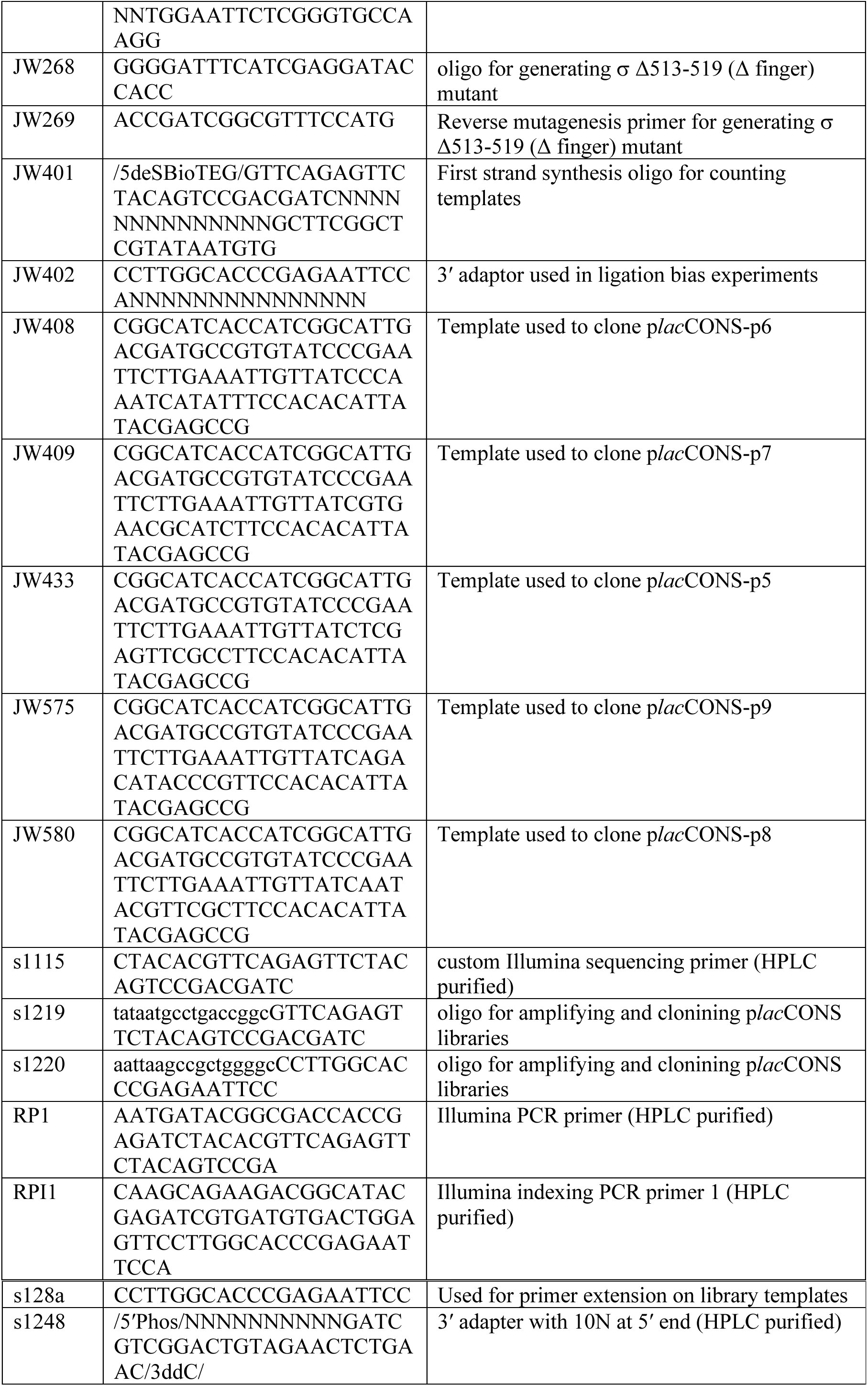
Oligonucleotides.

